# ACTIN Anchors the Highly Oligomeric DRP1 at Mitochondria-Sarcoplasmic Reticulum Contact Sites in Adult Murine Heart: Its Functional Implication

**DOI:** 10.1101/2021.11.29.470468

**Authors:** Celia Fernandez-Sanz, Sergio De La Fuente, Zuzana Nichtova, Yuexing Yuan, Sebastian Lanvermann, Hui-Ying Tsai, Marilen Ferderico, Yanguo Xin, Gyorgy Csordas, Wang Wang, Arnaud Mourier, Shey-Shing Sheu

**Affiliations:** Center for Translational Medicine, Department of Medicine, Sidney Kimmel Medical College, Thomas Jefferson University, Philadelphia, PA 19107 USA; Univ. Bordeaux, EPST, IBGC UMR 5095, 33077 Bordeaux Cedex, France; CNRS, IBGC CNRS UMR 5095, 33077 Bordeaux Cedex, France; MitoCare Center for Mitochondrial Imaging Research and Diagnostics, Department of Pathology, Anatomy & Cell Biol., Thomas Jefferson University, Philadelphia, PA 19107 USA; Mitochondria and Metabolism Center, Department of Anesthesiology and Pain Medicine, University of Washington, Seattle, WA 98109 USA

**Keywords:** Dynamin related protein 1 (DRP1), Mitochondria-associated membranes (MAM), β-ACTIN, Sarcoplasmic reticulum (SR), High Ca^2+^ microdomains, DRP1 oligomers, Adult cardiomyocytes

## Abstract

**Rationale:** Mitochondrial fission and fusion are relatively infrequent in adult cardiomyocytes compared to another cell types^1–3^. This is surprising considering that proteins involved in mitochondrial dynamics are highly expressed in the heart. It has been previously reported that dynamin-related protein 1 (DRP1) has a critical role in mitochondrial fitness and cardiac protection^1, 4^. Cardiac DRP1 ablation in the adult heart evokes a progressive dilated cardiac myopathy and lethal heart failure^1^. Nevertheless, the conditional cardiac-specific DRP1 knock-out animals present a significantly longer survival rate compared with global DRP1 KO models^1, 4, 5^. We have described before the great importance for cardiac physiology of the strategic positioning of mitochondrial proteins in the cardiac tissue^6, 7^. Therefore, we hypothesize that DRP1 plays a regulatory role in cardiac physiology and mitochondrial fitness by preferentially accumulating at mitochondria and junctional sarcoplasmic reticulum (jSR) contact sites, where the high Ca^2+^ microdomain is formed during excitation-contraction (EC) coupling.

**Objective:** This study aims to determine whether mitochondria-associated DRP1 is preferentially accumulated in the mitochondria and jSR contact sites, the mechanism responsible for such a biased distribution, and its functional implication.

**Methods and Results:** Using high-resolution imaging approaches, we found that mitochondria-associated DRP1 in cardiomyocytes was localized in the discrete regions where T-tubule, jSR, and mitochondria are adjacent to each other. Western blot results showed that mitochondria-bound DRP1 was restricted to the mitochondria-associated membranes (MAM), with undetectable levels in purified mitochondria. Furthermore, in comparison to the cytosolic DRP1, the membrane-bound DRP1 in SR and MAM fractions formed high molecular weight oligomers demosntratd by 2D blue native technique. In both electrically paced adult cardiomyocytes and Langendorff-perfused beating hearts, the oscillatory Ca^2+^ pulses preserved MAM-associated DRP1 accumulation. Interestingly, similar to DRP1, all mitochondria-bound β-ACTIN only exists in MAM and not in the purified mitochondria.

Additionally, co-immunoprecipitation pulls down both DRP1 and β-ACTIN together. Inhibition of β-ACTIN polymerization with Cytochalasin D disrupts the tight association between DRP1 and β-ACTIN. In cardiac-specific DRP1 knock-out mouse after 6 weeks of tamoxifen induction (DRP1icKo), the cardiomyocytes show disarray of sarcomere, a decrease of cardiac contraction, loss of mitochondrial membrane potential, significantly decreased spare respiratory capacity, and frequent occurrence of early after contraction (EAC), suggesting the heart is susceptible to arrhythmias and heart failure. Despite of this phenotype, DRP1icKo animals have longer life span than other DRP1 KO models. Strikingly, DRP1 levels are only modestly decreased in the MAM when compared with the rest of the cellular fractions. These preserved levels were accompanied by the preservation of the mitochondrial pool in the MAM fraction obtained from the DRP1icKO hearts.

**Conclusions:** The results show that in adult cardiomyocytes, mitochondria bound DRP1 clusters in high molecular weight protein complexes at MAM. This clustering is fortified by EC coupling mediated Ca^2+^ transients and requires its interaction with β-ACTIN. Together with the better preserved DRP1 levels in the DRP1icKO model in the MAM, we conclude that DRP1 is anchored at the mitochondria-SR interface through β-ACTIN and positions itself to play a fundamental role in regulating mitochondrial quality control in the working heart.

## INTRODUCTION

Mitochondrial dynamics, including fission, fusion, and motility, are fundamental mechanisms in regulating mitochondrial network morphology and mitochondrial subcellular distribution^8^. Mitochondrial dynamics also control several cellular processes, including cell division, oocyte maturation^8, 9^, axonal projection^10, 11^, autophagy, and apoptosis^12, 13^. Defective mitochondrial dynamics have been linked to neurodegenerative diseases, diabetes, cardiomyopathy, cancer, and several other human diseases^7–9^.

Mitochondrial dynamics are controlled by GTP hydrolyzing proteins, including dynamin-related protein 1 (DRP1) for fission and mitofusin 1 (MFN1), mitofusin 2 (MFN2), and optic atrophy 1 (OPA1) for fusion^17, 18^. In contrast to MFN1 and MFN2, which are physically anchored to the outer-mitochondrial membrane (OMM), DRP1 is a cytosolic protein. DRP1 undergoes GTP-cycle-dependent conformational changes to drive self-assembly and disassembly, but the structural bases of this process are still incompletely understood^19^.

DRP1 is detected ubiquitously in mammalian tissues, with the highest levels found in energy-demanding tissues like the heart, skeletal muscles, and brain^20, 21^. The high level of DRP1 in the heart seems to be at odds with numerous studies showing that mitochondrial dynamics in adult ventricular myocytes occur rather infrequently compared to another cell types^1–3, 22, 23^. Despite the scarcity of mitochondrial dynamics in adult cardiac mitochondria^24^, DRP1 is essential for controlling several heart functions being critical in heart development, suggesting a possible non-canonical role of DRP1 in the cardiac tissue^25–27^.

Mitochondria occupy ∼35% of the cell volume in the heart^28^. The majority of these mitochondria are located between the myofibers and appear fragmented with little mobility, anchored through β-ACTIN cytoskeleton architecture^29^ with a significant number tethered to the junctional sarcoplasmic reticulum (jSR) near the dyads^30^. The juxtaposition of mitochondria to the jSR leads to the formation of a high Ca^2+^ microdomain between the organelle contact areas when Ca^2+^ is released from the SR during excitation-contraction (EC) coupling^30^. This high Ca^2+^ microdomain provides privileged Ca^2+^ transport from jSR to the mitochondria through the mitochondrial Ca^2+^ uniporter complex (MCUC). It is widely believed that the mitochondrial Ca^2+^ uptake through MCUC stimulates the ATP regeneration processes rapidly and robustly, to meet the extremely high energy requirements for perpetual blood pumping, known as the excitation-bioenergetics (EB) coupling^31^. However, this concept has been challenged by recent findings showing that MCU knock-out causes little energy crisis in the beating heart^32–34^.

We recently reported a study showing that inhibition of DRP1 activity pharmacologically or genetically resulted in only a mild change in mitochondrial morphology in face of a significant decrease in State 3 respiration in adult cardiomyocytes^35^. These results uncover a novel non-canonical function of DRP1 in positively stimulating mitochondrial bioenergetics in adult cardiomyocyte, which is likely independent of mitochondrial fission. Interestingly, it has been reported that 97% of oligomerized DRP1 colocalizes with the mitochondria does not directly participate in fission events in cultured cells^36, 37^. We and others have shown that increases in cytosolic Ca^2+^ concentrations ([Ca^2+^]_c_) promote DRP1 translocation from Cytosol to mitochondria in cardiomyocytes^38^. Based upon these reported data, we hypothesize that in adult cardiomyocytes DRP1 is strategically localized in the mitochondria-jSR contact sites where high [Ca^2+^] microdomains are created during EC coupling. The unique clustering of DRP1 in this discrete area serves to regulate mitochondrial fitness and quality control sustaining the energetic balance of supply and demand in beating heart. To test this hypothesis, we have applied multiple techniques encompassing mouse echocardiography, mitochondrial respiration assays, IonOptix for measuring calcium dynamics and related sarcomere shortening, Airy-scan confocal imaging, and immunogold electron microscopy, sub-cellular fractionations, blue native polyacrylamide gel electrophoresis (BN-PAGE) and DRP1 inducible cardiac-specific knock-out (DRP1icKO) mouse model^1,4,39^.

The results showed for the first time, that in the heart, the mitochondrial membrane-associated DRP1 localized in high molecular weight clusters at the jSR-mitochondria interface, with a surprisingly no detectable DRP1 in the highly purified mitochondria membrane fraction. This preferential accumulation was preserved by mitochondria-associated β-ACTIN and EC coupling-linked Ca^2+^ transients. This unique feature of DRP1 oligomers positioning at the high Ca^2+^ functional microdomains between jSR and mitochondria could enable its non-canonical role in the heart.

## METHODS

### Animals

Experimental preparations were derived from the heart of 12- to 14-week-old male C57BL/6 mice, male Sprague-Dawley rats, and DRP1 inducible cardiac-specific knock-out (DRP1icKO) mouse^1, 4, 39^. The DRP1icKO was obtained by crossing the DRP1^fl/fl^ mouse (Dr. Junichi Sadoshima, Rutgers New Jersey Medical School) with the αMHC-Mer-Cre-Mer (Cre+) mouse line (JAX). Cre-mediated DRP1 ablation was induced by tamoxifen IP injection (40mg/kg/day) for 3 consecutive days. Sacrifice was made 6 weeks after the first injection. Reaming DRP1 levels were checked for Western blot from isolated cardiomyocyte lysate and by immunofluorescence in isolated cardiomyocytes from DRP1icKO (S1 A).

Animal handling was done in accordance with the National Institutes of Health Guide for the care and use of laboratory animals, and the protocols were applied in compliance with the Thomas Jefferson University institutional review board guidelines.

### Echocardiographic analysis

Transthoracic echocardiography was performed and analyzed on a Vevo2100 imaging system (VisualSonics. Inc., Toronto, Canada) to evaluate the changes in cardiac function. B-mode and M-mode images were collected in long and mid-papillary level short-axis orientations. All echocardiographic parameters were calculated by VevoLab 3.2.0 software. AI (Artificial Intelligence) based Auto-LV technology was used in the functional and anatomical analysis of the left ventricle to avoid inter-operator variability.

### Isolation of mouse cardiomyocytes

To obtain adult cardiomyocytes, 12- to 14-week-old C57BL/6 mice were sacrifice by cervical dislocation and anticoagulated using an IP heparin (50 IU/mice) injection. Hearts were retro-perfused through the aorta in a Langendorff system (Radnoti, Covina, CA) according to the protocol previously described^6, 40^. Freshly isolated adult cardiomyocytes were used in all experimental procedures.

### Electrical field stimulation on isolated adult cardiomyocytes

The electrical field stimulation experiments performed to elucidate DRP1 localization in adult cardiomyocytes were done as described before^6, 41^. Briefly, isolated calcium-tolerant cardiomyocytes from adult mice hearts were placed in 25 mm diameter coverslips pretreated with laminin and mounted in an electrical stimulation chamber equipped with two Pt electrodes (RC-47FSLP, Warner Instruments). Cells were continuously perfused with field stimulation buffer that contained in mM: 150 NaCl, 5.4 KCl, 10 HEPES acid, 1 glucose, 2 pyruvate, 5 creatine, and 5 taurine, supplemented with 2.5 μM or 2 mM CaCl_2_. Electrical biphasic pulses of 25-35 V amplitude, 2 Hz frequency, and 5 ms duration were applied to the cells for 15 min (MyoPacer Stimulator, IonOptix). Then, cardiomyocytes were fixed and subjected to a standard IF staining protocol for further analysis. To perform Ca^2+^ and contractility measurements, FURA2-AM loaded cardiomyocytes were monitor under IonOptix system (IonOptix LLC) paced at 1 Hz for 1 min followed by 30 sec resting period and 5 Hz for a minute. Cytosolic Ca^2+^ transients, early after contraction events (EAC) and sarcomere length, were registered and analyzed by IonWizard 6.6 software.

### Oxygen Consumption Rate (OCR) measurements from freshly isolated mouse cardiomyocytes

For measuring OCR in intact cardiomyocytes, we used the XF2e4 Extracellular Flux Analyzer (Seahorse Bioscience, Agilent Technologies). Equivalent sub-saturating densities of cardiomyocytes were plated on laminin-coated 24 well plates (10µg/µl). These sub-saturating densities allowed to reach linear responses of OCR. When measuring OCR, DMEM with 25 mM glucose and 0.1 mM pyruvate was used and 4 µM oligomycin A combined with 1 µM carboxyatractyloside (CATR), 1 µM carbonyl cyanide-p-trifluoromethoxyphenyl hydrazone (FCCP), and 10 µM antimycin A were added in three sequential injections^42, 43^. Data were normalized by citrate synthase activity measured by colorimetric assays^44^.

### Isolation of subcellular fractions from murine heart

Cytosol (Cyt), SR, crude mitochondria (cMit), pure mitochondria (pMit), and mitochondria-associated membranes (MAM) enriched fractions were obtained from murine heart homogenates by fractionation as described before^41^. Briefly, fresh mouse ventricles were minced and homogenized using a Potter–Elvehjem PTFE pestle-glass tube (loose followed by tight), on ice-cold isolation buffer containing (mM): 10 HEPES acid, 225 mannitol, 75 sucrose, 0.1 EGTA, pH = 7.4. After an initial centrifugation step (750 × g for 5 min, 4°C) the supernatant was centrifuged (7,000 × g for 10 min, 4°C). The resulting pellet contained crude mitochondrial fraction. The supernatant was centrifuged (40,000 × g for 45 min, 4°C) to obtain the SR-enriched fraction in the pellet. This pellet consists of SR that has small debris of functional mitochondria that remain attached to it as described before^6^. This fraction will be defined throughout this paper as SR. The supernatant was further ultracentrifuged (100,000 × g for 1 h, 4°C) to obtain the Cytosolic fraction in the supernatant. Crude mitochondrial fraction previously obtained was further purified in a 30% Percoll gradient ultracentrifuged (65,000 × g, for 30 min, 4°C) in a swing-out rotor, resulting in pure mitochondria (heavy fraction) and MAM (light fraction). Supplementary figure 2 (S2) shows a detailed schematic representation of the cardiac cellular fractions. The high content of RyR2 in the MAM fraction confirmed the extraction of jSR associated mitochondria.

### Protein analysis and Western blot

Equal amounts of protein (70 μg) supplemented with 5x Protein Loading Buffer (National Diagnostics, USA) were preheated (95°C, 5 min), separated on 12% acrylamide/bis (Bio-Rad) gels, and transferred to Amersham Hybond-ECL nitrocellulose membranes 0.2 µm (GE Healthcare). After blocking with Odyssey™ Blocking buffer (PBS) for 1 h at room temperature, the membranes were incubated with the primary antibodies of interest— DRP1 (BD; 61113), β-ACTIN (Santa Cruz Biotechnology Inc.; sc-4778), SERCA2 (Thermo Fisher; MA3-919), HSP60 (Cell Signaling Technology; 12165), ANT (Mitosciences; MSA02), TOM20 (Santa Cruz Biotechnology Inc.; sc-11415), Calsequestrin (Abcam; ab3516), FIS1 (ENZO; ALX-210), MiD49 (Proteintech; 16413-1-AP), MiD51 (Proteintech; 20164-1-AP), MFF (Abcam; ab129075), INF-2 (GeneTex; CTX130714), SPIRE-1C (custom made, Dr. Manor) TIM23 (BD; 611222), GAPDH (Cell Signaling Technology; 2118L) —in Odyssey™ Blocking buffer (PBS) overnight at 4°C. Infrared secondary antibodies IRDye 680LT goat anti-mouse and IRDye 800CW goat anti-rabbit (LI-COR Biosciences) and/or Horseradish Peroxidase (HRP)-conjugated goat-antimouse and goat-antirabbit antibodies (Cell Signaling Technology) were used for visualization of the proteins by LI-COR Odyssey scanner (LI-COR Biosciences). The specificity of the DRP1 antibody was checked for WB in isolated cardiomyocyte lysate from DRP1icKO (S1 A, right panel). Image Studio™ Lite software (LI-COR Biosciences) was used for band quantification and densitometry analysis.

### First dimension light blue native polyacrylamide gel electrophoresis (LBN-PAGE)

For LBN-PAGE, 100-300 μg of protein from each fraction of study were treated with extraction buffer pH 7.4 (10 μl/100μg) containing (mM): 30 HEPES, 150 CH3CO2K, 2 ε-amino-n-caproic acid, 1 EDTA, 12% (v/v) glycerol supplemented with 3% (w/v) digitonin (Sigma Aldrich) for 30 min at 4°C. Extracts were mixed with loading dye (1/400 of 5% [w/v]) Coomassie Brilliant Blue G-250, 150 mM BIS-TRIS, and 500 mM ε-amino-n-caproic acid, pH 7.0. Native PAGE Novex 3%–12% Bis-Tris Protein Gels (Life Technologies) were loaded with 100-300 μg extracts. After electrophoresis, lanes were stained with Coomassie Brilliant Blue R-250 or used for second dimension analysis.

### Second dimension SDS-PAGE

Lanes from the first-dimension Light Blue Native gels were first treated with equilibration buffer containing (mM): 6 Urea, 50 Tris-HCl, 30% glycerol, 2% SDS for 5 min. On a second step, lanes were treated with Equilibration Buffer supplemented with 200 mM DTT (Sigma Aldrich) for 30 min followed by treating with Equilibration Buffer supplemented with 135 mM Iodoacetamide (Sigma Aldrich) for 15 min. The remaining DTT and iodoacetamide were removed by a final washing step with Equilibration Buffer for 15 min.

Treated lanes were mounted on 4%-12% Bis-Tris NuPAGE™ Protein Gels (Novex™ Invitrogen™) and transferred to Amersham Hybond-ECL nitrocellulose membranes (GE Healthcare). The membranes were blocked in TBS-T solution (Tris-buffered saline, 0.1% Tween-20) with 5% non-fat milk powder. Proteins were detected with DRP1 in TBS-T with 3% non-fat milk powder. HRP-conjugated IgG anti-mouse antibodies (Cell Signaling Technology) were used as secondary antibodies. Peroxidase reactions were carried out and visualized using Supersignal West Dura Extended Duration Substrate (Thermo Scientific) and the chemiluminescence imaging system GBOX-Chemi-XRQ gel documentation system, Syngene. Image Studio™ Lite software (LI-COR Biosciences) was used for band quantification and densitometry analysis.

### Immunofluorescence (IF) and imaging analysis

Isolated cardiomyocytes were plated on laminin-coated coverslips and fixed (4% PFA). A 3% BSA-0.2% Triton®X-100 solution was used for permeabilization and blocking followed by a 1 h RT incubation with PBS–1% BSA containing mouse monoclonal DRP1 antibody (611113, BD Transduction Laboratories, 1:50) followed by a 1 h RT incubation with PBS–1%BSA containing rabbit polyclonal RyR2 (PA5-36121, Thermo Fisher, 1:50) and/or rabbit polyclonal TOM20 antibody (Santa Cruz, 1:50). Alexa Fluor®488 or 647 were used as secondary antibodies. SlowFade® was used for mounting on microscope slides. Specificity of the DRP1 antibody was checked for IF in DRP1KO-MEF cell line, adult cardiomyocytes from DRP1ciKO mice and rat adult cardiomyocytes treated with DRP1shRNA (S1 A-C).

Immunofluorescence was imaged using the Zeiss LSM880 system equipped with the Airyscan super-resolution (1.7x beyond diffraction limit) detection system. A 63X Zeiss plan-apochromat oil, 1.4 NA, DIC lens was used to obtain all images. Image analyses were done using FIJI/ImageJ (NIH).

For the RyR2/TOM20-DRP1 images, the same threshold was applied to each channel. Specifically, for the DRP1 immunolabeled images, a threshold considering only the 5% from the maximal intensity was applied. With this setting, only the brighter signals were considered, which is coincident with the DRP1 aggregates (puncta pattern). By using this mask, the resulting coincident particles from the two channels with > 0.05 μm^2^ area were counted. On a second step, the RyR2 signal was dilated by 66 nm and 132 nm (1 and 2 pixels) from the original labeling. By using this new mask, the same particle coincidence analysis strategy was applied. Five to ten microscope fields of 1275 µm^2^ were analyzed for each condition.

For the TOM20-DRP1 images, the same threshold was applied to each channel. By applying the same analysis strategy, the DRP1 particles found in contact with the outer mitochondria membrane (OMM) at different distances were counted. On a second step, the particles were classified by their localization at the transversal side (T side) or longitudinal side (L side) of the mitochondria for each condition. Supplementary figure 3 (S3) shows a detailed example of this analysis.

### Pre-embedding immunogold and transmission electron microscopy

Isolated mouse cardiomyocytes were fixed with 5% paraformaldehyde/PBS followed by quenching with glycine and permeabilization and first blocking with 3% BSA+0.2% Triton-X. Cardiomyocytes were incubated for 1 h in primary antibody diluted (1:100) in 1% BSA + 0.2% Triton-X, followed by secondary blocking with 5% goat serum in PBS. As secondary antibody FluoroNanogold™ (anti-mouse, 1:100) was diluted in 1% goat serum-0.1% TritonX-100 for 1 h. Primary antibody control for IG-TEM was performed in the absence of primary antibody, using the same secondary binding protocol as specified above (S1 C). Nanogold particles were developed for 3 min using GoldEnhance™ (Nanoprobes). Gold enhancement was followed by fixation with 1.6% glutaraldehyde and 0.2% tannic acid in PBS; post-fixation with 1% osmium tetroxide in 0.1M sodium cacodylate buffer; and contrasting with 1% aqueous uranyl acetate. The cells were dehydrated in graded alcohol and embedded in Durcupan. Upon polymerization, the coverglass was dissolved using concentrated hydrofluoric acid, after which ultrathin sections (65-80 nm) were prepared using a Leica UTC ultramicrotome (w/Diatome diamond knife) and imaged with a thermos/FEI transmission electron microscope (Tecnai 12, Hillsboro, OR, USA). Images were taken at 3,200-15,000x magnification (at 80 keV). Analysis of gold particles was performed using FIJI/ImageJ (NIH).

Mitochondrial morphology assessment was performed as described before^7^. For the analysis of the DRP1-particles distribution around the mitochondria, the T side and L side were considered as the minor and major axes, respectively, and compared with the total perimeter of the organelle (S4).

### Langendorff perfusion of rat hearts

Adult male Sprague–Dawley rats (250–300 g) were anesthetized with isoflurane (3% in oxygen) and submitted to a bilateral thoracotomy. Whole hearts were quickly excised and retrogradely perfused through the aorta with an oxygenated (95% O_2_–5% CO_2_) Krebs solution at 37°C in mM: 118 NaCl, 4.7 KCl, 1.2 MgSO_4_, 25 NaHCO_3_, 1.2 KH_2_PO_4_, and 11 glucose, pH 7.4 in a constant-flow Langendorff system, as previously described^45^. After 15 min of equilibration, two hearts were treated in normoxic conditions with low or physiological concentrations of Ca^2+^ (2.5 µM or 1.8 mM respectively). Each concentration was applied for 60 min. After the treatments, hearts were collected, and different cellular fractions were obtained as described above.

### Statistical analysis

Data are expressed as mean ± S.E.M., and two-tailed Student’s t-test was used for comparisons between two independent sample groups. Factorial ANOVA analysis was applied for comparison of groups with more than one factor, by *a priory* comparison when necessary. Differences of p <0.05 were considered statistically significant (two-tailed significance was commonly applied and one-tailed when justified by previous data and predictions about the direction of the difference). When samples did not follow a normal distribution, the non-parametric test of the median was used. All statistical analyses were performed with IBM-SPSS v.24 software (New York, NY, USA).

## RESULTS

### DRP1 is preferentially located at the SR-mitochondria interface in primary adult cardiomyocytes-

In order to test our hypothesis that DRP1 may preferentially accumulate at SR and mitochondria junctions nearby the high Ca^2+^ microdomains created by Ca^2+^ release from SR during EC coupling we proceed to characterize the subcellular distribution of DRP1 by using multiple imaging techniques. Immunofluorescence (IF) localization of DRP1 was obtained by using the Airyscan detector of the Zeiss LSM880 confocal system (Fig. 1 A, C). This high-resolution imaging system allowed us to quantitatively evaluate the distance between the location of DRP1 with respect to the location of mitochondria (labeled by TOM20) (Fig. 1 A) and jSR (labeled by RyR2) (Fig. 1 C). The pixel size in the collected images is 66X66 nm square. Therefore, we measured the radiated distances from DRP1 fluorescent punctate particles to that of TOM20 or RyR2 in 66 nm increments (66 nm, 132 nm, and 198 nm). When the distance measured between DRP1 puncta and TOM20 or RyR2 puncta was < 66nm, then the two puncta were located within the same pixel and thus, we considered them as colocalization (overlap). When the distance was < 132 nm, then the two proteins were located in the next adjacent pixel, or 2 pixels away if the distance was < 198 nm. Similar imaging analysis was previously used by us to show that the mitochondrial Ca^2+^ uniporter (MCU) was predominately localized toward mitochondria-jSR contact sites where RyR2 is located^6^. Quantitative analysis shows that 83.76% ± 2.60 of the DRP1 punctate particles (likely representing oligomerized DRP1 because the lower background fluorescent puncta were removed through thresholding) were overlapped with TOM20 fluorescent particles, and 93.95 ± 2.06% were located in the next adjacent pixel (<132nm) (Fig. 1 B). These results were consistent with previous observations in various cell lines (e.g., human osteosarcoma U2OS cells), showing that the majority of DRP1 puncta are associated with mitochondria^36, 37^. For DRP1 and RyR2 puncta, the same analysis strategy revealed that 71.02% ± 3.40 of DRP1 particles were located within 196 nm radius of RyR2 fluorescent particles (Fig. 1 D), suggesting that DRP1 oligomers are accumulated at the proximity of the jSR. Note that the longer distances between DRP1 and RyR2 than DRP1 and TOM20 were mostly due to that RyR2 are predominantly localized in jSR facing the T-tubule (plasma membrane) and not facing mitochondria as we have previously reported^30^.

**Figure 1.**
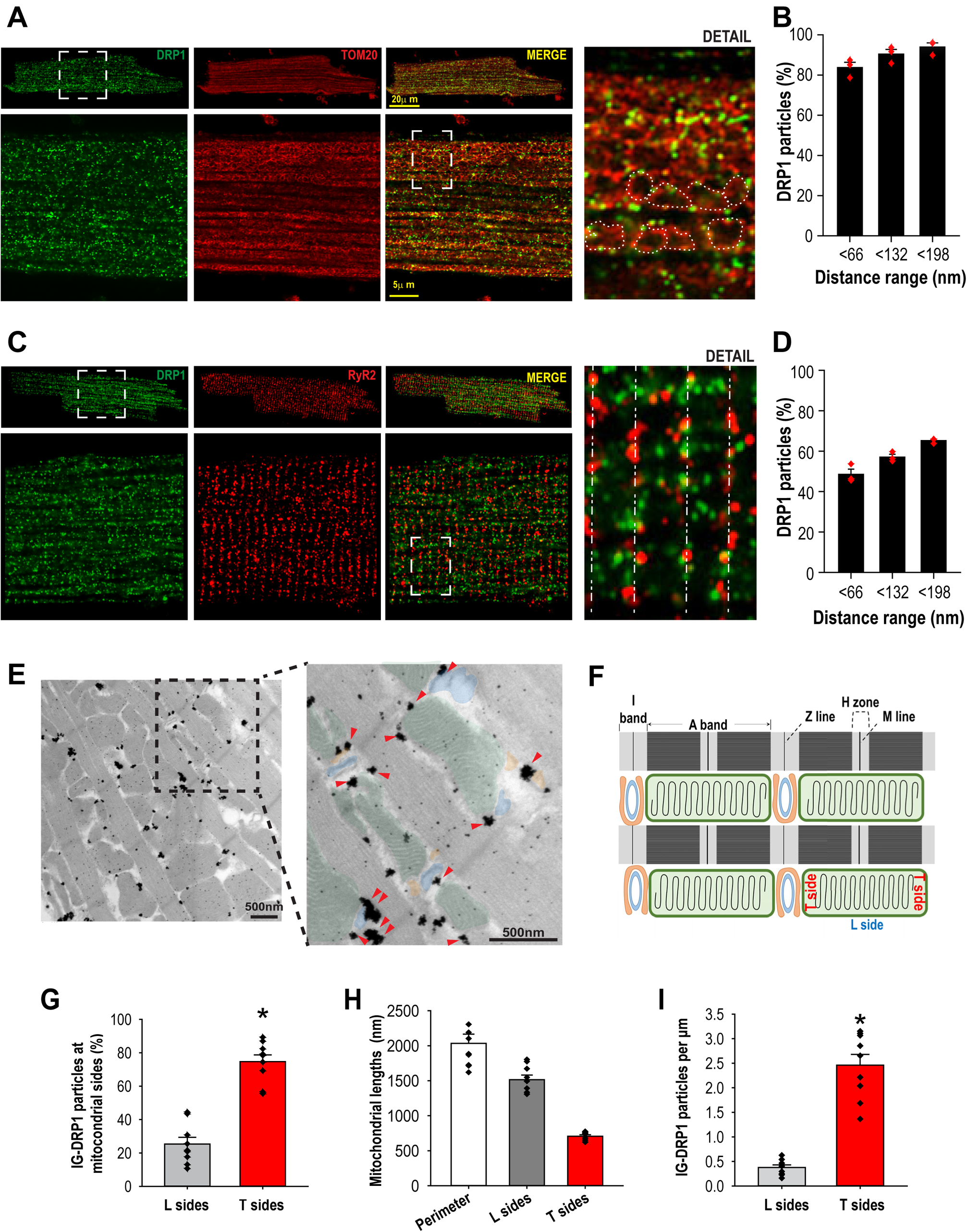
DRP1 is preferentially located at SR-mitochondria interface in adult Cardiomyocytes. **A)** Confocal images showing immunostaining of DRP1 (green), TOM20 (red), and merging of the two images in adult cardiac myocytes. Lower panels: zoomed images adapted from the dashed line box after AiryScan processing. “Detail” zoomed image adapted from the dashed line bracket in the adjacent panel demonstrates the proximity of DRP1 puncta with mitochondrial OMM (encircled by the white dots). **B**) Summarized data showing the fraction of DRP1 within the indicated distance ranges (<66, <132 and <198 nm) from TOM20. N=3 animals, 7-9 cells per animal. **C)** Confocal images showing immunostaining of DRP1 (green) and RyR2 (red) in an adult cardiac myocyte. Lower panels: zoomed images are after AiryScan processing. “Detail” zoomed image at far-right panel shows the proximity of DRP1 with RyR2 along the Z-lines (dashed line). **D)** Summarized data showing the fraction of DRP1 within the indicated distance ranges (<66, <132 and <198 nm) from RyR2. N=3 animals, 7-9 cells per animal. **E)** Immunogold TEM detection of DRP1 in adult cardiac myocytes. Electron microscopy of DRP1 labeled with Immunogold (IG). Zoom: colored labels highlight SR (orange), T-tubules (blue), and mitochondria (green). Red arrows indicate DRP1 particles. **F)** For quantitative analysis of the location of DRP1-IG particles, a schematic diagram is generated to illustrate the transversal (T) side of mitochondria where jSR, T tubule, and mitochondria are in juxtaposition and longitudinal (L) side of mitochondria where no jSR is in proximity. **G)** Summarized data showing the percentage of DRP1-IG positive particles associated with the L or T side of the mitochondria. **H)** the lengths of the mitochondrial perimeter, L sides and T sides (nm). N= 4 cells, 2-4 fields per cell, 122 mitochondria, ***: p<0.001. **I)** Representation of the IG-DRP1 particle per µm was obtained by normalizing the number of particles per side (T or L) by the length of each side (µm). N= 4 cells, 2-4 fields per cell, total of 115 mitochondria, ***: p<0.001.

We next used pre-embedding immunogold TEM labeling of DRP1 (DRP1-IG) to reveal its subcellular localization at higher resolution. For an easier view, the key structural organelles are color-coded; orange for the SR, blue for the T-tubule, and green for the mitochondria. As highlighted by the zoomed area with red arrows pointing at DRP1-IG particles, the larger size particles appeared to locate more abundantly in the area where mitochondria, SR and T-tubule were in proximity (Fig. 1 E). These DRP1 IG-positive particles are specific in labeling DRP1 as they were significantly diminished in the no-primary control and DRP1icKO samples (S1 D). For a better understanding of the quantitative analysis of the location of DRP1-IG particles at the different regions of mitochondria, a schematic diagram was created to illustrate the transversal side of mitochondria (T) where jSR, T tubule, and mitochondria are in juxtaposition and longitudinal side of mitochondria (L) where jSR is not in proximity (Fig. 1 F, S4). The quantitative analysis of DRP1-IG particles in the T and L sides is summarized in Fig. 1 G, which shows 74% ± 4.05 of the particles on the T side. On average, the T sides represent ∼1/3 while L sides represent ∼2/3 of the total mitochondrial perimeter (Fig. 1 H). Therefore, when normalized to the length, the number of particles per µm in T side was 8.09 ± 1.68-fold higher than that in the L side (Fig. 1 I). Altogether, these data show that DRP1 is strategically localized at the jSR-mitochondria contacts, in proximity to the Ca^2+^ release units formed by the RyR2 on the SR and the L-type Ca^2+^ channels on the T-tubules (dyads)^30^.

### Mitochondria SR contacts harbor the membrane bound DRP1-

The observation from high-resolution microscopy and IG-TEM indicated that DRP1 preferentially localizes at the jSR-mitochondria interface and prompted us to further determine the association between DRP1 and SR-mitochondrial membranes. To this end, subcellular fractions of heart ventricles were analyzed by Western blot. Our results indicate that DRP1 is differently distributed among the Cytosolic (Cyt), SR, and crude mitochondrial (cMit) fractions (Fig. 2 A). The Cytosolic fraction was validated by the absence of SR/ER Ca^2+^ ATPase (SERCA), calsequestrin (CSQ), the mitochondrial import receptor subunit (TOM20), and adenine nucleotide translocase (ANT); the SR fraction was validated by the higher abundance of SERCA and CSQ and the lower abundance of TOM20 and ANT; and the cMit fraction was validated by the higher abundance of TOM20 and ANT and the lower abundance of SERCA and CSQ. These characteristics of each cardiac subcellular fraction are consistent with our previously reported subfractionations^6^. Note that the cardiac SR is a structurally diverse organelle that consists of junctional (RyR2 and CSQ enriched) and network (SERCA-enriched) SR (jSR nd nSR respectively)^46^. Unfortunately, no biochemical method can jSR and nSR, therefore the SR subfraction in this study was a mixture of the two components. Furthermore, it has been more notably appreciated in recent years that SR/ER and mitochondria are tightly tethered by several proteins^47–49^ and that SR/ER can “wrap” around the mitochondria^50^, resulting in a logical cross “contamination” between the SR and mitochondrial subfraction. As indicated by the SR fraction contains mitochondrial proteins and vice versa. This phenomenon is specific to cardiac tissue since when we compared SR/ER fractions from the heart and liver, no traces of mitochondria were detected in the latter (S 5). Based on these criteria, the quantitative comparison of the relative amount of DRP1, SERCA, CSQ, TOM20, and ANT, between the Cytosolic *vs.* the SR and cMit fractions were plotted (Fig. 2 B, C). The analysis showed that DRP1 was 2.25 ± 0.23-fold higher in the Cytosol than in SR (Fig. 2 B) and 8.05 ± 1.72-fold higher in the Cytosol than in cMit (Fig. 2 D). Surprisingly, when comparing the SR *vs.* cMit fraction, DRP1 was 3.64 ± 0.26-fold higher in the SR than in the cMit fraction (Fig. 2 D), suggesting a significant accumulation of DRP1 in the SR fraction.

**Figure 2.**
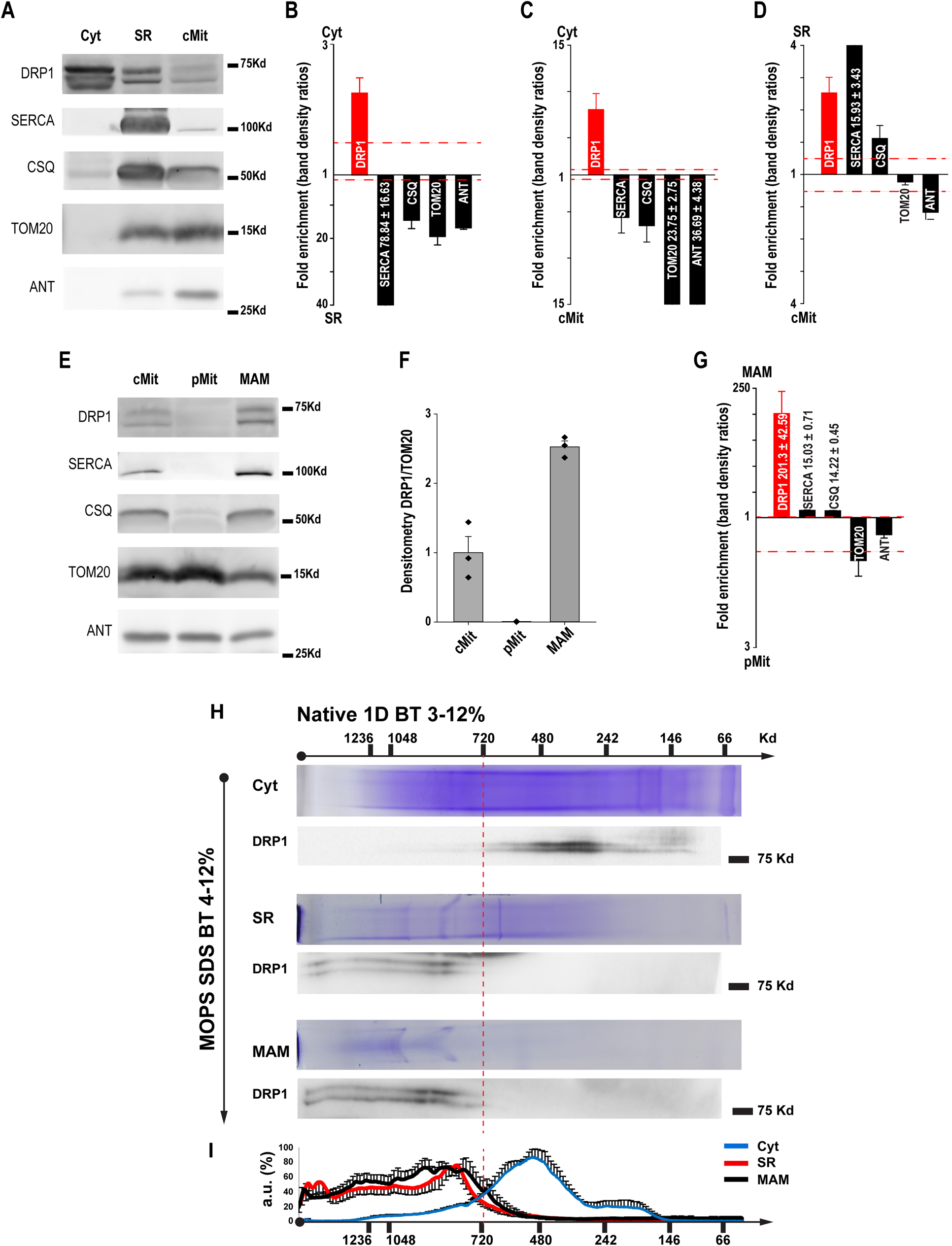
DRP1 is localized at the SR and MAM fractions in high molecular weight oligomers. **A)** Representative Western blot of the cardiac subcellular fractions obtained by differential centrifugation, Cytosol (Cyt), sarcoplasmic reticulum (SR), and crude mitochondria (cMit). The sarcoplasmic/endoplasmic proteins reticulum ATPase (SERCA) and calsequestrin (CSQ) were used as SR markers while mitochondrial import receptor subunit (TOM20) and adenine nucleotide translocase (ANT) were used as outer and inner mitochondrial membrane markers respectively (OMM and IMM). **B-D)** Comparisons of relative abundance of proteins between the Cyt, SR and cMit fractions. **E)** Representative Western blot of the mitochondrial fractions obtained by Percoll® purification. The same protein markers as in panel A were used. **F)** Western blot analysis of the mitochondrial fractions showed a DRP1 enrichment in the mitochondria-SR associations (MAM) but not in the purified mitochondrial fraction (pMit). **G)** Comparison of relative abundance of proteins between the pMit and MAM fractions. N = 3 fractionations. **H)** Solubilized proteins from the Cytosol, MAM, and SR subcellular fractions were separated according to their masses on a linear 3%-12% acrylamide gradient gel for BN-PAGE (first dimension, 1D). Native protein complexes were separated by 4%-12% Bis-Tris SDS-PAGE gradient gel (second dimension, 2D). **I)** Densitometry analysis of the DRP1 oligomers distribution in the different fractions N = 3-5 fractionations.

The cMit fraction was then further purified using a Percoll® gradient separating pure mitochondria (pMit) from MAM (Fig. 2 E) to elucidate the DRP1 positioning within these two sub-mitochondrial regions. Remarkably, similar to the SR markers SERCA and CSQ, DRP1 was undetectable in the pMit fraction while practically all DRP1 from the cMit was recovered in the MAM fraction, which also contains both SR (SERCA and CSQ) and mitochondrial markers (TOM20 and ANT) (Fig. 2 E). The normalized densitometry of DRP1 indicated a high presence of DRP1 in the MAM, with no detectable traces of DRP1 in the pMit (2.52 ± 0.08 vs 0.13 ± 0.035 respectively, Fig. 2 F). The comparison of fold enrichments of DRP1, SERCA, CSQ, TOM20, and ANT between MAM and pMit was plotted in Fig. 2 G. The results showed that DRP1 was highly enriched (201.3 ± 42.6-fold enrichment) in the MAM vs the pMit fraction (Fig. 2 G).

These results show that, in cardiac tissue, the mitochondria-associated DRP1 exclusively localizes at the MAM while the specific mechanism that promotes DRP1 localization at the SR-mitochondrial interface within adult cardiomyocytes is still unknown.

### DRP1 localized at the SR and MAM fractions at high oligomeric State-

As described before, the main difference between the soluble and membrane-bound DRP1 resides in its oligomerization state^51^. Therefore, a comprehensive analysis to elucidate the oligomerization states of the DRP1 in the different cellular subfractions was performed. This same experimental approach would exclude the possibility of cross-contamination of this soluble form into the membranous fractions (SR and MAM). To this end, DRP1 complexes from the different fractions were solubilized using digitonin and resolved by Blue Native-PAGE. Solubilized proteins from the Cytosol, SR, and MAM fractions were separated according to their size on a linear 3%-12% Blue Native (BN)-PAGE gradient gel (first dimension, 1D) (Fig. 2 H, Coomassie Blue lanes). Native protein complexes were then separated by 4%-12% Bis-Tris SDS-PAGE gradient gel (second dimension, 2D) (Fig. 2 A, blots). The densitometry analysis of the 2D experiments revealed that the SR and MAM fractions contained a similar pattern, with higher molecular weight DRP1 complexes, in comparison with the smaller DRP1 complexes found in the Cytosolic fraction (Fig. 2 I). No DRP1 was detected on the 2D experiments from purified mitochondria (S6). The non-overlapping oligomeric nature of the DRP1 in the soluble fraction *vs*. the membranous fractions confirms that high DRP1 oligomers always appear attached to membranous fractions, while small DRP1 oligomers remain soluble in the Cytosol. Additionally, the experiment ruled out the possibility of cross-contamination between the Cytosol and membranous fractions.

Overall, these data, together with the IF and DRP1-IG results, indicate that highly polymerized DRP1 is preferentially localized at the SR and MAM but not in the Cytosol.

### Ca^2+^ transient activity is involved in DRP1 positioning at the SR-mitochondria associations in the heart

As shown before, DRP1 plays a critical role in controlling cardiac bioenergetics. It has been extensively reported that mitochondrial Ca^2+^ signaling also contributes significantly to regulate cardiac bioenergetics. Therefore, the high Ca^2+^ microdomains created at the SR-mitochondrial junctions could be a key factor implicated in the DRP1 accumulation at these functional microdomains. To test this idea, cardiomyocytes were isolated and acutely submitted to 15 min-2 Hz electrical field stimulation while being continuously perfused with Ca^2+^ (2 mM) or quasi-Ca^2+^-free (2.5 μM) extracellular buffer. Cells were immediately fixed and immunolabeled for DRP1 and TOM20 (S7) or RyR2 (Fig. 3 A) or immunolabeled for IG-TEM (Fig. 3 C). In the IF images, the distribution of DRP1 particles aggregates in relation to the L and T sides of the mitochondria (TOM20, red) was analyzed. The results showed a significantly higher amount of DRP1 oligomers at the T sides of the mitochondria when cardiomyocytes were stimulated in the presence of 2 mM Ca^2+^ in comparison with those stimulated with 2.5 μM Ca^2+^ (S7). The same experimental strategy was performed to study the distribution of DRP1 particles in relation to the SR constituting the dyads (RyR2 labeling). The analysis of DRP1 distribution along the RyR2 from isolated cardiomyocytes revealed a significant amount of DRP1 particles at the RyR2 vicinity upon field stimulation in the presence of 2 mM Ca^2+^ (Fig. 3. B). In the area within 1 pixel distance (66 nm), there was 39.71% ± 2.00 of DRP1 colocalized with RyR2 in the presence of 2 mM Ca^2+^. This colocalization decreased to 26.08% ± 1.75 in the condition of 2.5 µM Ca^2+^, a 1.58-fold difference. In the area within 2 pixels distance (132 nm) or 3 pixels distance (192 nm), the difference of DRP1 and RyR2 colocalization between 2 mM and 2.5 µM Ca^2+^ became smaller gradually; at 132 nm the colocalization was 56.11% ± 2.67 in 2 mM Ca^2+^ and 41.97% ± 2.48 in 2,5 µM Ca^2+^, a 1.38-fold difference, and became even smaller at 192 nm (1.29-fold difference). This gradual decrease in the percentage of DRP1 and RyR2 colocalization as their separation distance becomes larger suggests that the Ca^2+^-mediated DRP1 accumulation in the vicinity of RyR2 is rooted to the very close SR-mitochondria contact sites, where the microdomains of Ca^2+^ concentrations are presumably highest.

**Figure 3.**
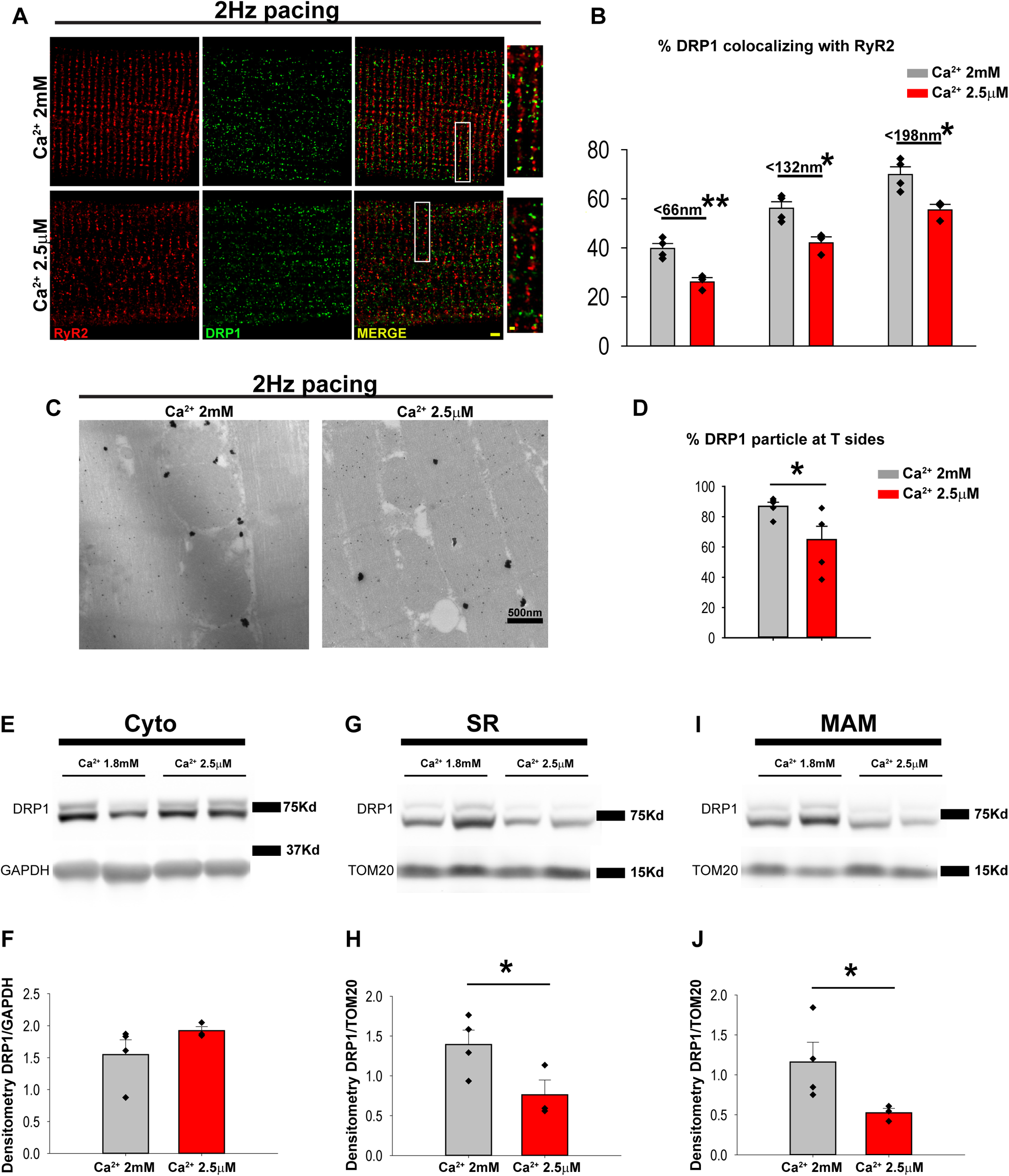
Ca^2+^ signaling takes part in DRP1 positioning at the cardiac SR-mitochondria contacts. Cardiomyocytes were isolated and submitted to electrical field stimulation (15 min-2 Hz) while being superfused Ca^2+^ (2 mM) or quasi-Ca^2+^-free (2.5 μM) buffer. **A)** Zoomed images (scale bar = 2 µm) and detail (scale bar = 0.5 µm) are after AiryScan processing DRP1 (green) and RyR2 (red). **B)** Summarized data showing the fraction of DRP1 within the indicated distance ranges (< 66, < 132, and < 198 nm) from RyR2. (N = 3 animals; 10 cells per animal) *: 0.05 > p > 0.01, **: 0.01 ≥ p ≥ 0.001. **C**) Immunogold-DRP1particles in field stimulated adult cardiac myocytes perfused with different [Ca^2+^]. **D)** Analysis of the IG-DRP1 particle distributions found at the mitochondrial T sides and Sub-mitochondrial fractions of beating rat heart perfused with either 1.8 mM Ca^2+^ buffer or quasi Ca^2+^ free (2.5 µM) for 60 min. Representative Western blot and quantification of DRP1 levels in the Cytosol **(E, F)**, SR **(G, H)** and MAM **(I, J).** N = 3-4 rats per condition. *: 0.05 > p > 0.01, one tailed t-test.

In order to reveal how Ca^2+^ it is involve in subcellular localization with a higher resolution, we performed IG-TEM analysis from field stimulated cardiomyocytes (Fig. 3 C). The obtained data showed a significant reduction of the DRP1 particles at the T side of the mitochondria in cardiomyocytes perfused with a quasi-Ca^2+^-free buffer in comparison with those perfused with physiological [Ca^2+^] (2 mM) (86.84% ± 2.72 *vs.* 64.83% ± 8.82, Fig. 3 D).

To confirm the previous results in whole heart preparations, spontaneously beating rat hearts were Langendorff-perfused with Krebs buffer supplemented with 2.5 µM or 1.8 mM Ca^2+^ for 60 min before being subjected to homogenization and fractionation. Western blot analysis of the fractions showed no changes in the amount of cytosolic DRP1 due to the different levels of Ca^2+^ in the perfusion buffer (Fig. 3 E, F). Interestingly, and consistently with the previous results, the levels of DRP1 were significantly lower in the SR and MAM fractions from the hearts treated with low Ca^2+^ buffer (1.39 ± 0.36 *vs.* 0.76 ± 0.18 for the SR and 1.16 ± 0.28 *vs*. 0.52 ± 0.05 for the MAM, Fig. 3 G-J). Despite the treatments with different Ca^2+^ concentrations, no DRP1 was detected in the pMit (S8).

Taken together, these results suggest that the high [Ca^2+^] at the dyads stabilizes high-molecular-weight DRP1 complexes positioned within the SR-mitochondria associations in the beating heart, preventing its delocalization.

### β-ACTIN is responsible for DRP1 accumulation in MAM

Besides Ca^2+^, DRP1 positioning at the mitochondrial membrane strictly depends on anchoring proteins^52–56^. We studied the distribution among the fractions of the previously described DRP1 anchoring proteins such as mitochondrial fission factor (MFF), mitochondrial fission 1 (FIS1), and mitochondrial dynamics proteins 49 and 51 (MiD49, MiD51)<u>. Western blot analysis showed that</u> <u>none of the described anchoring proteins</u> followed the characteristic DRP1 distribution among the subcellular fractions in the heart (Fig. 4 A, B) suggesting additional mechanisms may be responsible for anchoring DRP1 in MAM.

**Figure 4.**
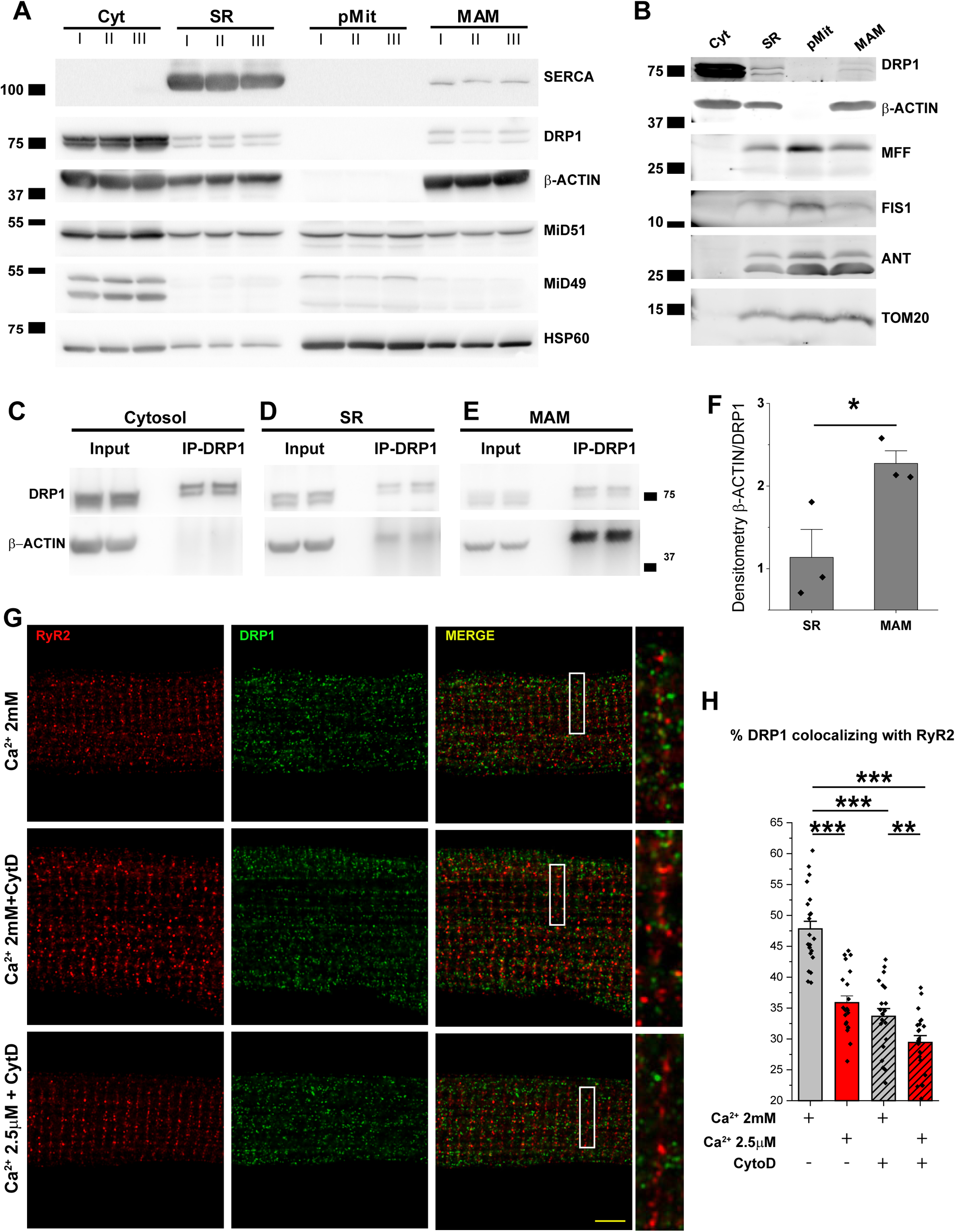
β-ACTIN co-precipitates with DRP1 at the mitochondria-SR associations and not in the Cytosol. **A-B)** Representative WB of DRP1 anchoring proteins – MFF, FIS1, MiD49 and MiD51-distribution among the different cellular fractions (Cyt, SR, pMit and MAM, N=5 animlas per fractionation, 3 different fractionations). **C-E**) DRP1-IP and β-ACTIN Co-IP for Cyt, Sr and MAM fractions. **F)** WB quantification of β-ACTIN coimmunoprecipitating with DRP1 in the SR and MAM fractions (N=5 animlas per fractionation, 3 different fractionations for each DRP1-IP, *: 0.05 > p > 0.01). **<u>G)</u>** Cardiomyocytes were isolated and pretreated with cytochalasin D 10μM for 30min in standard culture medium. Then the cells were submitted to electrical field stimulation (30 min-1 Hz) while being superfused Ca^2+^ (2 mM) or quasi-Ca^2+^-free (2.5 μM) buffer in the presence or absence of cytochalasin D 10μM. Zoomed images (scale bar = 5 µm) and detail (are after AiryScan processing DRP1 (green) and RyR2 (red). **H)** Summarized data showing the fraction of DRP1 within the indicated distance ranges (< 66nm) from RyR2. (N = 3 animals; 10 cells per animal) **: 0.01 ≥ p ≥ 0.001, ***: p<0.001.

It has been shown that β-ACTIN polymerization at or near fission sites can stimulate oligomeric maturation of DRP1 on mitochondria filaments and thus may serve as a dynamic reservoir for recruiting oligomerized DRP1 to the ER-mitochondria contact sites in osteosarcoma (U-2 OS) cells^57^. In addition, increases in intracellular Ca^2+^ concentrations induce β-ACTIN polymerization^58^.

We wanted to verify whether if there was a direct interaction between DRP1 and β-ACTIN, if this interaction was specific in the mitochondria-SR associations and the regulatory role of Ca^2+^ in this process. DRP1 was immunoprecipitated from Cytosolic, SR, and MAM fractions. We observed that, despite the high abundance of DRP1 and β-ACTIN in the Cytosol (Fig 4 C), β-ACTIN was only Co-precipitating with DRP1 in the membranous fractions, SR and MAM (Fig 4 D, E). Interestingly, a significantly higher specific interaction between DRP1 and β-ACTIN was present in the MAM when compared with the SR fraction (Fig 4 D-F), showing a significantly stronger binding between DRP1 and β-ACTIN at the mitochondria associated membranes. When inhibiting β-ACTIN by treating freshly isolated cardiomyocytes with Cytochalasin-D under pacing, we observed a DRP1 delocalization from the RyR2 vicinity. DRP1 diffusion from the RyR2 vicinity was exacerbated by combining the Cytochalasin-D treatment with low Ca^2+^ concentration (2.5 μM) (Fig. 4 G-H).

These data that support the hypothesis that β-ACTIN and Ca^2+^ are central players for the DRP1 accumulation in MAM.

### DRP1 knock-out leads to impaired contractility in the heart and isolated cardiomyocytes without significant changes in mitochondrial morphology

After elucidating the specific localization of DRP1 in murine cardiomyocytes, we decided to pursue the importance of DRP1 in regulating cardiac function in adult hearts. Therefore, we performed a series of physiological experiments to determine the cardiac phenotype in DRP1icKO mice. This model was chosen due to the embryonic lethality of the DRP1-null mice^59^. Similar to the previous report^1^, echocardiographic analysis revealed a severely reduced contractile dysfunction associated with DRP1 deficiency in the heart. Our results show that, compared with the controls, DRP1icKO mice have a significant decrease in cardiac functional parameters (ejection fraction Ctrl= 52.42% ± 1.92 *vs.* DRP1icKO= 26.98% ± 2.02, fractional shortening Ctrl= 26.68% ± 1.22 *vs*. DRP1icKO= 12.47% ± 1.00, cardiac output Ctrl= 15.99 ml/min ± 0.68 *vs.* DRP1icKO= 10.01 ml/min ± 0.77 and stroke volume Ctrl= 34.64µl ± 1.35 *vs*. DRP1icKO= 20.05 µl ± 1.31 (S9).

To elucidate the cellular mechanisms beneath the pathophysiological alterations in cardiac performance *in vivo*, we studied the consequences of DRP1 removal on the EC-coupling in single cardiomyocytes isolated from adult DRP1icKO mice. In the following experiments, isolated cardiomyocytes were submitted to field stimulation (1Hz and 5Hz) to measure cytosolic Ca^2+^ transients and cell contractility (isotonic contraction/cell shortening).

At 1 Hz, the dynamics of cytosolic Ca^2+^ transients were significantly altered, with a prolonged time to peak and to relaxation without changes in Ca^2+^ transients amplitude (Fig 5 A, B, S-table 1), mimicking the Ca^2+^ transients recorded from the cardiomyocytes of failing heart as reported previously^60^. The much longer Ca^2+^ transients subject the cardiomyocytes to a higher propensity of EAC as shown previously in failing heart^61^.

**Figure 5.**
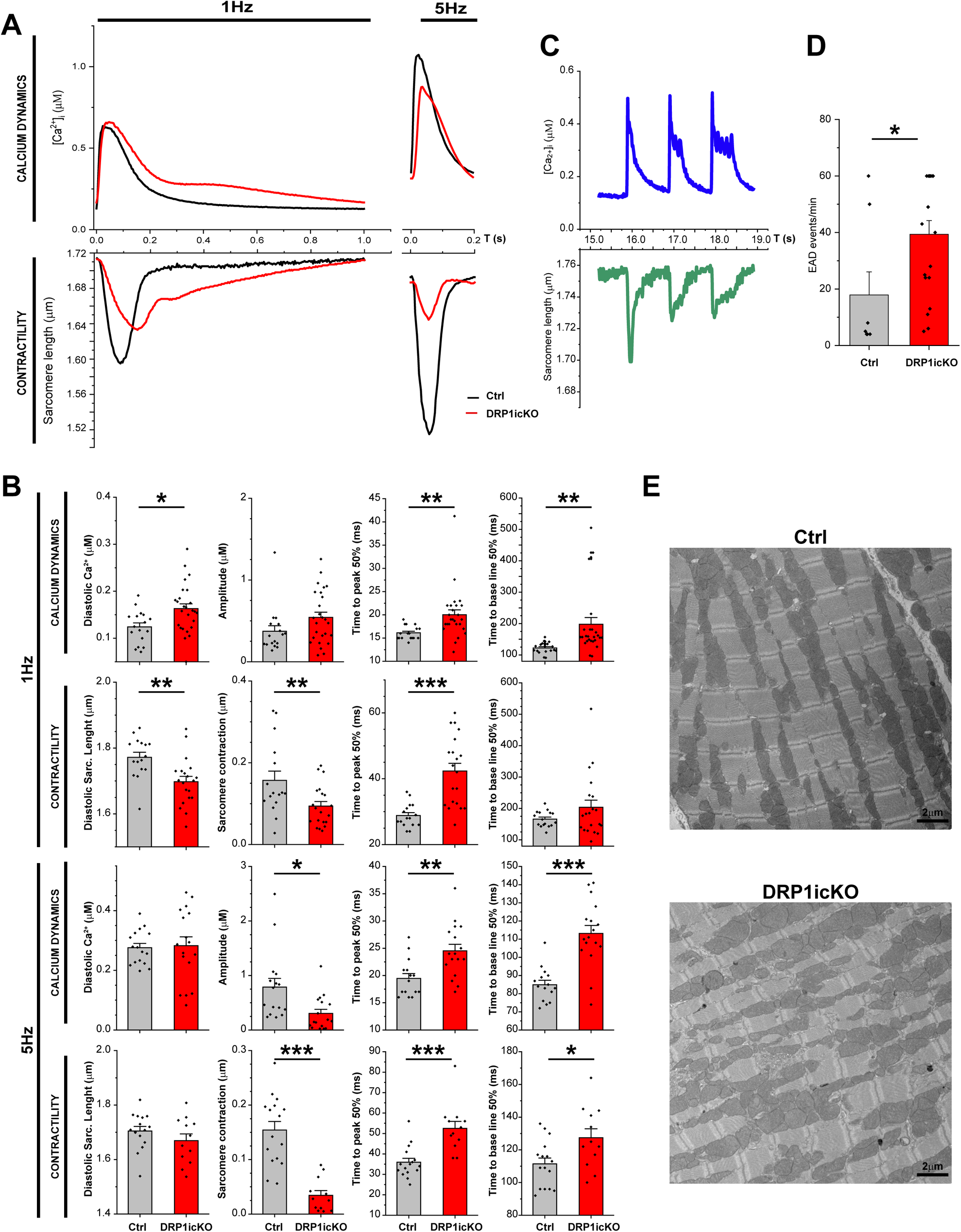
Ablation of DRP1 in the adult heart is related with altered Ca^2+^ dynamics, contractility and increased EAD and structural aberrancies. **A)** Representative traces of cytosolic calcium transients and sarcomere length for control and DRP1icKO cardiomyocytes were freshly isolated and submitted to electrical field stimulation (1.5 min at 1 Hz) and 5 Hz while being superfused in the presence of 100nm Isoproterenol. B) Bar graph representation of Ca^2+^ dynamics and sarcomere length parameters at 1Hz and 5Hz for control and DRP1icKO cardiomyocytes. N=3-4 animals per genotype, 6-10 cells per animal: *: 0.05 > p > 0.01, **: 0.01 ≥ p ≥ 0.001, ***: p<0.001. C) Representative traces of cytosolic calcium transients and early after depolarization events (EAD, black arrows). D) Bar chart showing the quantification of EAD events per minute detected in Ctrl and DRP1icKO isolated cardiomyocytes. N=3 independent experiments per genotype, 4 to 10 cells per experiment, *: 0.05 > p > 0.01. E) Transmitted electron microscopy images from Ctrl and DRP1icKO showing Z lines disarrangement caused by the depletion of DRP1.

Indeed, under the same experimental conditions, cardiomyocytes isolated from DRP1icKO mice showed a significantly higher frequency of spontaneous Ca^2+^ oscillations, indicative of EAC events when compared to the control counterparts (DRP1icKO= 39.37 EAC/min ± 4.87 *vs*. Ctrl= 17.88 EAC/min ± 8.18) (Fig 5 C, D). The higher incidents of EAC are consistent with the higher incidents of premature death of DRP1icKO mouse probably due to cardiac arrhythmias. At a higher stimulation frequency (5Hz), the Ca^2+^ dynamics impairment was accompanied by a decreased in the Ca^2+^ transient amplitude in the DRP1icKO (0.59 µM Ca^2+^± 0.09) cardiomyocytes compared to the controls (1.05 µM Ca^2+^± 0.17) (Fig. 5 A, B, S table 2).

A significantly diminished (∼40% at 1Hz and ∼70% at 5Hz) sarcomere shortening in cardiomyocytes isolated from the DRP1icKO was observed when compared with controls ( (Fig 5 A-B, S-table 3-4). The time to peak and time to recovery of contraction were significantly prolonged in DRP1icKO cardiomyocytes at both pacing frequencies (Fig 5 B, S table 3-4). These changes in single cardiomyocytes contraction can explain the decreases in the functional parameters of cardiac performance *in vivo* as described above. In addition to the impairments in cytosolic Ca^2+^ dynamics and contractions, severe Z-lines disarrangement was observed in DRP1icKO cardiomyocytes by TEM. This Z-lines disarrangement may contribute to the contractility impairment noticed in the DRP1icKO at organ and cellular level (Fig 5. E).

### Spare respiratory capacity and mitochondrial membrane potential are significantly reduced in DRP1 deficient adult cardiomyocytes

We previously published the results showing that pharmacological or genetic inhibition of DRP1 leads to a decrease in mitochondrial respiration with a minimal change in mitochondrial morphology^1^. It is well-known that the energy deficiency contributes to the loss of cardiac performance and contractility. Consequently, we measured mitochondrial oxygen consumption rate (OCR) in intact isolated cardiomyocytes from Ctrl and DRP1icKO mice (Fig 6 A) to evaluate its energetic stare. When compared with Ctrl, DRP1icKO presented a significantly increased basal respiration (Ctrl= 2472.4 pmol/ml*CS ± 500.4 vs DRP1icKO= 4765.4 ± 602.6) and significantly decreased spare respiratory capacity (Ctrl= 100% ± 15.1 vs DRP1icKO= 56.1% ± 9.4, % of the Ctrl) (Fig 6 B), indicating diastolic dysfunction and inefficiency in ATP production in response to increased energy demand. Consistent with the respiration measurements in intact cardiomyocytes, the mitochondrial membrane potential in DRP1icKO cardiomyocytes was significantly decreased (DRP1icKO= 348.534 a.u.f. ± 779.89 *vs.* Ctrl= 9059.52 a.u.f. ± 1154.27) (Fig. 6 C, D).

**Figure 6.**
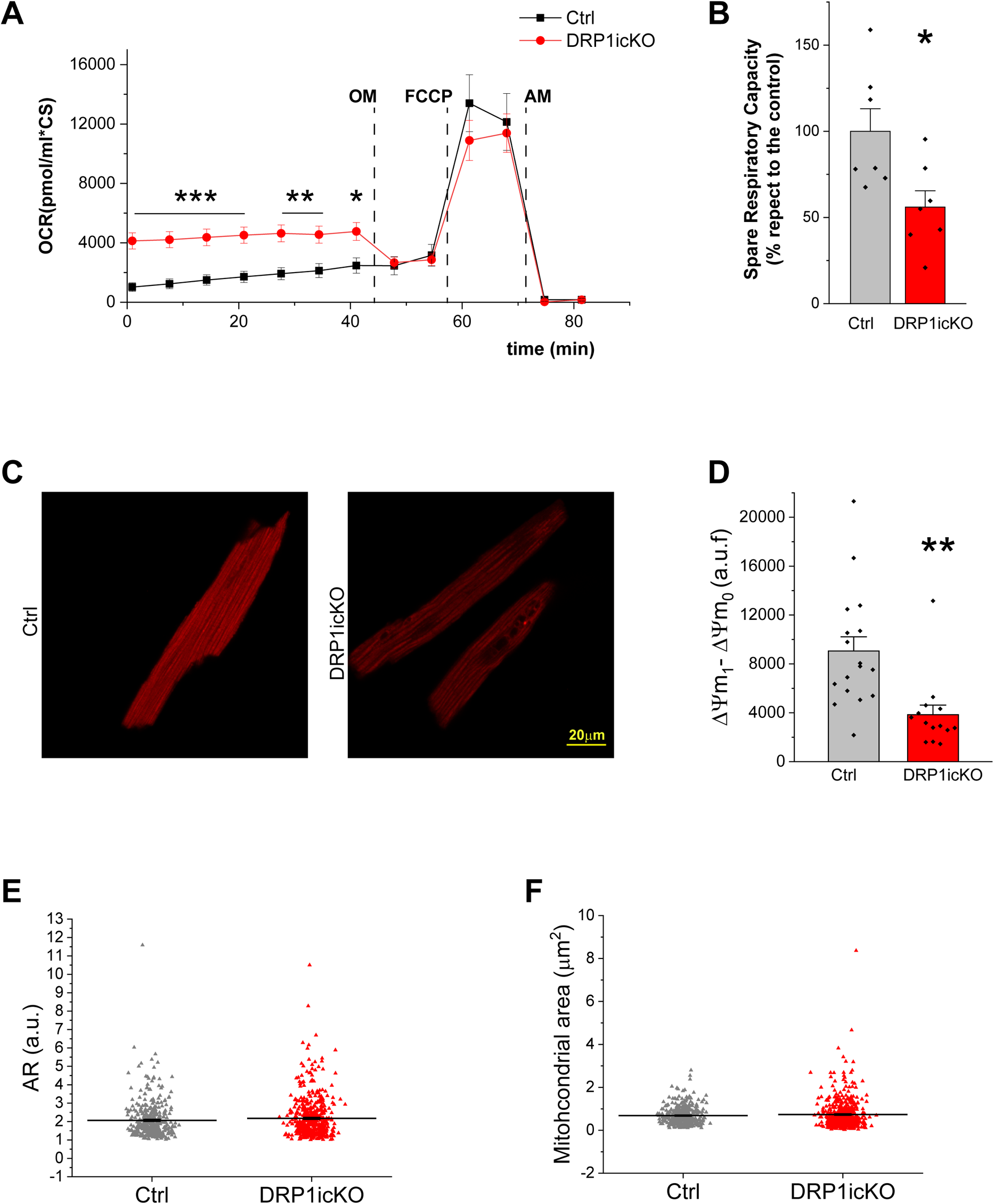
Isolated adult cardiomyocytes from DRP1icKO show a severally decreased spare respiratory capacity associated with a lower mitochondrial membrane potential. **A)** Metabolic profile and oxygen consumption rates (OCR) under basal conditions, after oligomycin (OM), FCCP and antimycin from control (black) and DRP1icKO (red) adult cardiomyocytes. B) Spare respiratory capacity was calculated by subtracting the basal OCR from maximal oxygen consumption obtained by the titration of exposure to FCCP and normalized to the control. n= 4 independent isolations per group, *: 0.05 > p > 0.01, **: 0.01 ≥ p ≥ 0.001. C) Representative images and D) bar chart showing the difference between the initial and final membrane potential of Ctrl and DRP1icKO, monitored by TMRE loading. Maximal depolarization was achieved by adding oligomycin (2μg/ml) and FCCP (μM). E-F) Mitochondrial aspect ratio and area quantification for Ctrl and DRP1icKO (N= 4 cells per group, 191 to 323 mitochondria per group)

A detailed analysis of the mitochondrial morphology from TEM images of DRP1icKO cardiomyocytes showed a normal-appearing mitochondria, with no significant changes in the mitochondrial aspect ratio (A.R.) or mitochondrial area (Fig. 6 E, F).

These results show that in the heart, DRP1 has a critical relevance in cardiac bioenergetics and mitochondrial fitness beyond its canonical role in mitochondrial fission.

### A significant portion of MAM associated β-ACTIN and DRP1 preserved in face of DRP1 ablation

As mentioned before, the DRP1 protein levels decreased 84.8% in DRP1icKO heart tissue when compared with control animals (S1, S10 A). Despite this significant DRP1 depletion, DRP1icKO animals are able to survive for at least 12 weeks^1, 4^. We analyzed the subcellular distribution of the remaining DRP1 at the different cellular fractions by western blot, from Ctrl and DRP1icKO mice. Only minimal traces of DRP1 were found in the Cytosolic fraction (8.7% respect to the Ctrl, S10 B). At the SR fraction, a decrease of 79.9% % was observed in the DRP1icKO animals. This diminished DRP1 expression was accompanied by a significant decrease of β-ACTIN of the same magnitude (Fig. 7 A-C). Interestingly, the decreases of DRP1 and β-ACTIN in the MAM fraction were much smaller (49%) in the DRP1icKO animals as if they try to adapt to the adverse conditions of DRP1 deletion. We also wanted to study the expression levels of the main proteins involved in the SR Ca^2+^ handling: RyR2 and SERCA. Following the tendencies of DRP1 and β-ACTIN, RyR2 and SERCA showed significantly reduced expression in the SR fraction from the DRP1icKO (∼50% for both proteins). In the MAM fraction, RyR2 levels were maintained while the SERCA levels showed a milder decrease when compared with the SR fraction changes (Fig. 7 D-E). We also observed that the mitochondrial pool and citrate synthase (CS) activity relative to the cardiac tissue (Fig. 7 F-G) was dramatically decreased in the pure mitochondrial fraction (72.2% less in the DRP1icKO vs. Ctrl) while it appeared unaltered in the MAM fractions were the DRP1 levels are better preserved.

**Figure 7.**
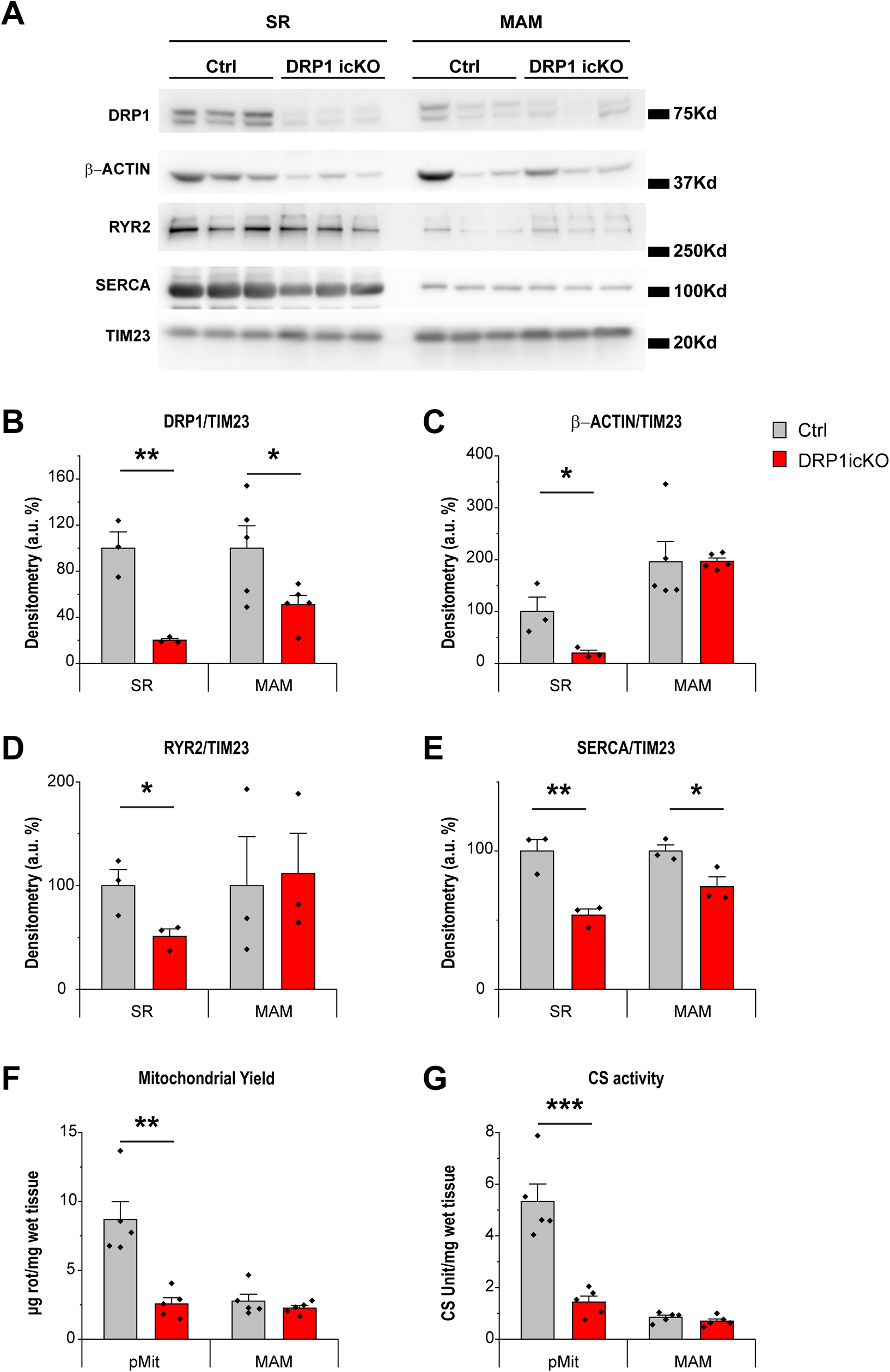
DRP1 levels are preserves/promotes at the MAM fraction in the DRP1icKO mice. **A)** WB analysis of SR and MAM fractions obtained from control and DRP1icKO animals after 6 weeks of tamoxiphen injection. **B-E)** Quantification of DRP1, β-ACTIN, RYR2 and SERCA levels in the SR and MAM fractions from both genotypes (N=5 animals per fractionation, 3 different fractionations per genotype, *: 0.05 > p > 0.01, **: 0.01 ≥ p ≥ 0.001). **F)** Mitochondrial yield and G) Citrate synthase Activity respect to wet tissue from pMit and MAM obtained from control and DRP1icKO animals after 6 weeks of tamoxiphen injection (N=5 animals per fractionation, 5 different fractionations per genotype, **: 0.01 ≥ p ≥ 0.001, ***: p<0.001).

This data, together with the IP results showed before, suggest that the strong interaction between β-ACTIN and DRP1 in the MAM helps to preserve its positioning at this fraction, even in face of DRP1 ablation. The unique feature of DRP1 oligomers positioning at the high Ca^2+^ functional microdomains between jSR and mitochondria, in strong association with β-ACTIN, could enable DRP1 non-canonical role regulating mitochondrial health in the heart, preserving cardiac function, increasing th lifespan of these animals.

## DISCUSSION

This study shows for the first time that, in adult murine cardiomyocytes under physiological conditions, DRP1forms high molecular weight oligomers and strategically localizes in the mitochondria-jSR associations but not in SR-free mitochondria. This unique pattern of localization is mediated by a strong association with β-ACTIN and regulated by Ca^2+^ from EC coupling. Genetic deletion of cardiac-specific DRP1 6 weeks after tamoxifen injection decreases the total cellular DRP1 by 84.8% (91.3% in the Cytosol, and 79.9% in the SR). Unexpectedly, DRP1 levels are better preserved in the MAM. The heart shows some degrees of functional deficit. However, the mice still appear to be healthy without any occurrence of death, noted that the global DRP1 KO is embryonic lethal^39^ and DRP1icKO die after 12 weeks of induction^1, 4^.

The tactical localization of other mitochondrial protein complexes involved in [Ca^2+^]_m_ dynamics and ECB coupling either clustered (MCU) or excluded (NCLX) from the SR-mitochondria interface has been described before by our goup^6, 7^. Similarly, here we report that DRP1 oligomers by their interaction with β-ACTIN and the EC coupling-associated Ca^2+^ transients, accumulate in this exquisite area^62^ (Fig. 8).

**Figure 8.**
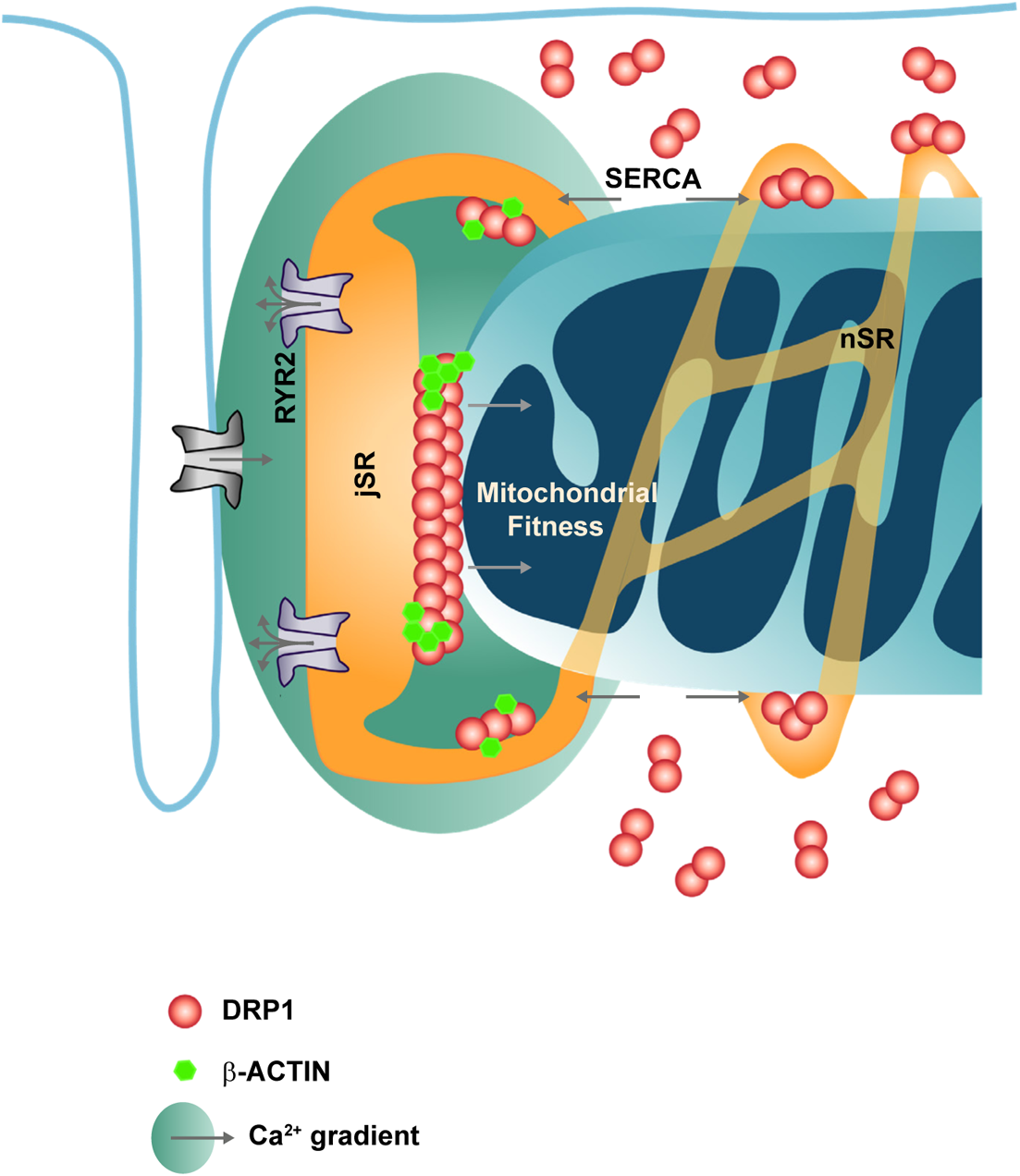
Mechanisms of high molecular weight DRP1 clusters at the SR and MAM in the beating heart. Schematic representation of the proposed mechanism by which Ca^2+^ signaling and β-ACTIN interaction play an essential role in DRP1 oligomerization and localization in the mitochondrial-associated membranes (see text for detailed description).

Our immunofluorescence study using Airyscan confocal microscopy and IG-TEM to obtain 2-dimensional images of DRP1 distributions showed that in adult cardiomyocytes, approximately 75% of mitochondria-associated DRP1 oligomers are found in close proximity to the areas that contain jSR (T side) (Fig. 1 C-I). The structural and functional microdomains between jSR and mitochondria are the hubs not only for a privileged Ca^2+^ exchange^30^ but also for other key cellular activities, such as redox signaling, lipid transfer, and mitochondria quality control^63–65^. The clustering of high molecular weight DRP1 protein complexes in this narrow space may intervene in the close interaction between these two organelles, enabling them to perform these crucial cell functions.

A detailed study on the DRP1 distribution among different cellular fractions of the heart revealed that DRP1 is preferentially localized at the SR-mitochondrial associations and not in the pure mitochondria, in close proximity to the Ca^2+^ release units formed by the RyR2 on the SR and the L-type Ca^2+^ channels on the T-tubules (Fig. 2 A-G, S 11). We have observed that SR fraction contains mostly the SR membrane with a smaller amount of tether mitochondrial membranes based on the quantitation of each organelle’s resident proteins (Fig. 2 A, D), phenomenon typically observed in the cardiac tissue (S5). The existence of DRP1 in ER has been observed previously. An earlier study has suggested that ER-localized DRP1 participates in the formation of nascent secretory vesicles from the ER cisternae^66^. In line with this observation, recent studies in cell lines have reported that some of the DRP1 anchoring proteins are found not only in the mitochondria but also at the ER surface^2, 37, 67^. In human osteosarcoma U2OS cells, ER can function as a platform for DRP1 oligomerization and transferring to mitochondria for mitochondrial fission^2, 36, 37^. Furthermore, β-ACTIN oligomerization enhances ER-mitochondria interaction allowing a more efficient Ca^2+^ transport from ER to mitochondria through mitochondrial Ca^2+^ uniporter (MCU)^3^. As described before by others, the main difference between the soluble and membrane-bound DRP1 rests on its oligomerization state^51, 68^. In addition to high-resolution imaging and WB measurements to show the DRP1 unique distribution at the SR-mitochondrial junctions, our results from Blue Native Page gels further demonstrate that in the heart, higher DRP1 complexes are found in the SR and MAM fractions. In comparison, significantly smaller complexes are present in the Cytosolic fraction. Since most of the cellular DRP1 is located in the Cytosol, experiments also ruled out the possibility that the membranous SR and MAM fractions were contaminated with the cytosolic form of DRP1 (Fig. 2 H-I).

The detailed mechanism responsible for the preferential accumulation of DRP1 at the jSR mitochondria associations still remains elusive. The findings in this study show that the SR and especially the MAM fractions are enriched in the proteins forming the Ca^2+^ release units —the RyR2 and L-type Ca^2+^ channel (S11)— suggesting that the high Ca^2+^ concentrations in these microdomains may facilitate DRP1 accumulation in the SR-mitochondria microdomains. Accordingly, our results show that Ca^2+^ dynamics of the beating heart are crucial for the DRP1 positioning at this jSR-mitochondria interface (Fig 3).

Besides Ca^2+^ transients as part of the DRP1 positioning mechanism, we studied the distribution of the most relevant DRP1 anchoring proteins to elucidate whether they follow the same pattern tan DRP1, acting as a landing platform to recruit DRP1 to jSR-mitochondria contacts. It has been previously described that DRP1 assembles to the OMM in a GTP binding, hydrolysis, and nucleotide exchanging-dependent manner^52^ and by interacting with adaptors and anchoring proteins such as Mff, FIS1, and MiD49/MiD51^53–56^. Our Western blot results from subcellular fractions demonstrate that none of the previously described anchoring proteins follow DRP1 distribution (Fig 4 A-B). Most of the anchoring proteins appear to be enriched in the pMit fraction, where DRP1 is undetectable. This data is consistent with previous observations where most of the proposed adaptors are evenly distributed over the mitochondria rather than specifically localized in MAM^54, 56^ or form weak interactions and might not be essential for the DRP1 assembly to the mitochondrial surface^53, 55^. This, together with our data, suggest that these anchoring proteins might be implicated in docking functions, but they are not defining the final positioning of DRP1 at the SR-mitochondrial contacts in adult cardiomyocytes.

From the DRP1 anchoring proteins study, we observed that only β-ACTIN appears to distribute as DRP1 among the cellular fractions (Fig 4 A-B). We also demonstrate that the distribution of both proteins is due to a strong and direct interaction between DRP1 and β-ACTIN at the MAM fraction (Fig. 4 C-F).

Ca^2+^ is an important regulatory mechanism on β-ACTIN polymerization^69^. Our results show that the Ca^2+^ dependent DRP1 localization described above is impaired by β-ACTIN depolymerization strategies (Fig 4 G-H). These results indicate that cytosolic Ca^2+^ transients in the beating heart are a regulatory mechanism of the β-ACTIN dependent DRP1 positioning at the SR-mitochondrial contact. Recent reports propose that, in cell culture models, β-ACTIN and the β-ACTIN-associated proteins, like INF2 and SPIRE1C, play a pivotal role in DRP1 recruitment to the nucleation sites in ER and mitochondria, respectively, with Ca^2+^ playing a critical role in the process^2, 3, 23, 36, 37^. How these proteins trigger, ER-mitochondrial interaction remains unclear, especially in the heart, where these β-ACTIN-interacting proteins do not distribute among the fractions as DRP1 and β-ACTIN (S12).

DRP1 has an essential role in regulating cardiac physiology at the organ and cellular level without impact on mitochondrial morphology (Fig. 5–6). These results are in accordance with previous studies where DRP1 depletion leads to contractile dysfunction and cardiac fibrosis, with minimal changes in mitochondrial dynamics^1, 35^. Additionally, we have observed that depletion of DRP1 leads to Z line misalignment. Misalignment of Z lines in myofibrils (Fig. 5 E), as well as some other factors like energy deficiency (Fig. 6), often lead to desynchronization of the calcium signal spreading^70, 71^, explaining the decreased cell contraction and increased frequency of EAC observed in the DRP1icKO cardiomyocytes under pacing (Fig. 5).

The loss of cardiac performance and contractility is frequently related to bioenergetics deficiencies. The obtained data from mitochondrial respiration analysis show that cardiomyocytes from control animals present modest basal respiration. This is a common observation in quiescent non-working cardiomyocytes that hydrolyze very little ATP and consequently consume little O_2_^42^. Cardiomyocytes from DRPicKO hearts present a significantly increased basal respiration (Fig. 6). This increased ATP consumption may be in part due to the diastolic dysfunction in DRPicKO that shows a small but significant increase in resting tension (shortening, Fig 5 A, B, S-table 1-4) as well as other energy-requiring pathological processes associated with the DRP1 depletion, like for example, increased mitophagy^1^. This increased basal respiration is associated with a significantly decreased ΔΨm and a consequent reduction in the spare respiratory capacity (Fig. 6), reflecting a significantly decreased capability of the DRP1icKO cells to produce extra ATP by oxidative phosphorylation in case of a sudden increase in energy demand but still compatible with life. DRP1 depletion is related to embryonic and postnatal lethality. Nevertheless, ablation of DRP1 in the adult heart is accompanied by a prolonged survival rate^1, 4, 5, 39^, probably due to compensatory mechanisms.

Our physiological data from the DRP1icKo mice, along with the observation from others^1^, show a progressive cardiac decompensation ending in lethal heart failure. DRP1 depletion appears to be responsible for a great mitochondrial loss due to an increase mitophagy in which underlies the progressive but tolerated cardiac dysfunction^1, 5^. Strikingly we show for the first time that when we KO *Drp1* in the adult heart, the DRP1 and β-ACTIN levels in the MAM fraction is better preserved, while the levels of DRP1 in the rest of the fractions are dramtically decreased. These stable DRP1 levels at the MAM are mirrored by the stabilizations of RyR2 and SERCA2A levels at these microdomains (Fig. 7 A-D). Additionally, we observed how mitochondrial yield is preserved in the MAM fraction of the DRP1icKO while the pure mitochondrial pool decreases by more than 70% (Fig 7 F-G). This preserved mitochondrial fitness has been described in the crude mitochondrial fraction from the DRP1icKO model before, reflected by a normal respiratory function and absence of increased ROS^1^. These observations coincide with our results since most of the crude mitochondria of the DRP1icKO are in its majority MAM due to the loss of the pure mitochondrial fraction.

We introduce in this study the novel concept that, in adult cardiomyocytes, the membrane-associated DRP1 forms very high molecular weight (up to millions of daltons) complexes in association with β-ACTIN and is strategically localized at jSR-mitochondria contact sites (Fig. 8). During excitation-contraction cycles, the oscillatory high [Ca^2+^] in jSR-mitochondria interfaces promote DRP1 oligomerization and accumulation at the SR-mitochondria interface. This process appears not to lead to fission events^24^ or changes in DRP1 phosphorylation under physiological conditions^72, 73^. Together with our previously published results^3, 35, 38, 74^, we suggest that, in the beating heart, physiological [Ca^2+^] and DRP1/β-ACTIN interaction is critical for the optimal DRP1 positioning at the jSR-mitochondria interface, preserving the cardiac structure and mitochondrial fitness. In the DRP1icKO model, the DRP1 better-held levels in the MAM help maintain mitochondrial fitness, contributing to the observed prolonged lifespan of these animals compared to other DRP1KO animal models.

This study also opens several questions that will require further investigation. In particular, if so, which post-translational modifications are involved in DRP1 strategic positioning under physiological [Ca^2+^]_c_? What are the specific proteins of the high DRP1 molecular weight complex (DRP1-β-ACTIN interactome) critical for its necessary arrangement in the heart? Finally, it would be of great interest to investigate the possible role of the DRP1-β-ACTIN complex on mitochondrial bioenergetics regulation in cardiac tissue.

## ACKNOWLEDGMENTS

We thank Ms. Jennifer Wilson for the English language editing on the manuscript. We also thank the lab members of Drs. Sheu, Mourier, Wang, Csordas for valuable discussions. Special thanks to Dr. Junichi Sadoshima for providing DRP1icKO mice, Dr. Uri Manor for supplying the custom-made Spire1C antibody, and Dr. Hiromi Sesaki for sharing the *Drp1* ko cell line. We thank Dr. Jyotsna Mishra for her contribution in setting up the DRP1 IP. We would also like to thank Dr. Rojo, and Dr. Duvezin-Caubet for their support and valuable technical and theoretical inputs.

## AUTHOR CONTRIBUTIONS

C.F-S., designed, performed, and analyzed the experiments with the technical assistance of S.L and H. T. S.DFP., participated in the field stimulation experiments, IF and analysis, and contributed to the writing and discussion of the manuscript. Z.N., performed and participated in the analysis of the EM and IG studies. C.F-S., wrote the first draft of the manuscript. G.C., W.W., A.M., and S-S.S., oversaw the whole project.

## SOURCES OF FOUNDING

This work was supported by NIH fundings: HL137266 to S-S.S., and W.W. and HL142864 to SS.S., and G.C., and partially supported by the American Heart Association (18EIA33900041 to W. W.). A.M., receives support from the AFM-Telethon Trampoline (AFM-19613) and ANR (ANR-16-CE14-0013). C.F-S., received funding from the French State in the framework of the “Investments for the Future” Program, IdEx Bordeaux, reference “ANR-10-IDEX-03-02”, and from the American Heart Association - 2020 AHA Postdoctoral Fellowship, reference “20POST35211149”.

