## Supplementary Figures for "ACTIN Anchors the Highly Oligomeric DRP1 at Mitochondria-Sarcoplasmic Reticulum Contact Sites in Adult Murine Heart: Its Functional Implication" (from another version of this preprint: version 2, 2021-12-02)

**Supplementary figure 1**

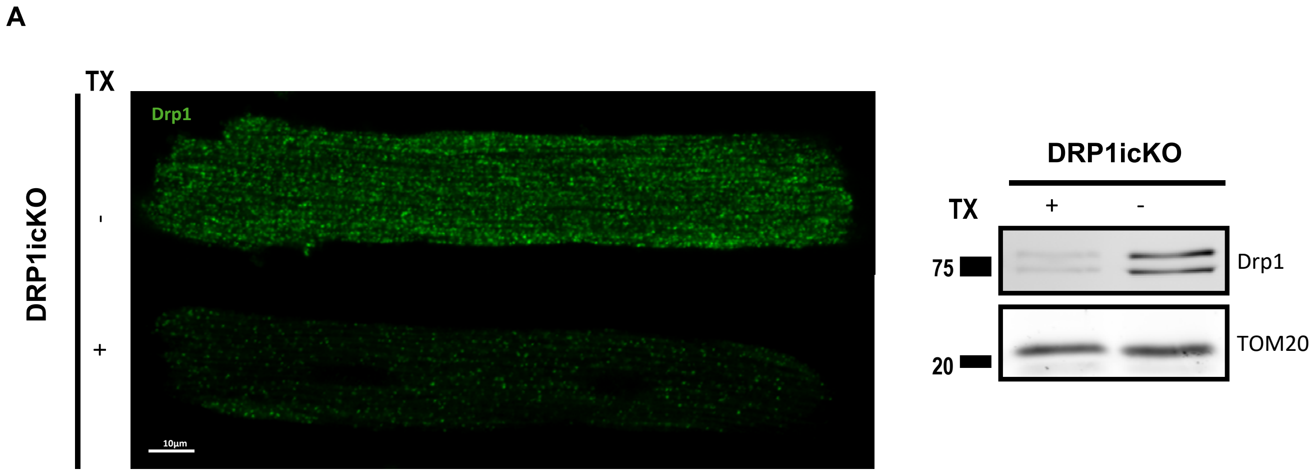

**Fig. S1A. Verification of DRP1 antibody and ciKODRP1 mice in IF and WB from freshly isolated cardiomyocytes lysate.** Anti-DRP1 antibody (BD Biosciences, 611112) detects mitochondrial associated punctate DRP1 “hot spots” in wild type but not DRP1cKO cardiomyocytes after 4 weeks tamoxifen induction (left panel). On the right, WB analysis from cardiomyocytes lysate from control and tamoxifen induced mice.

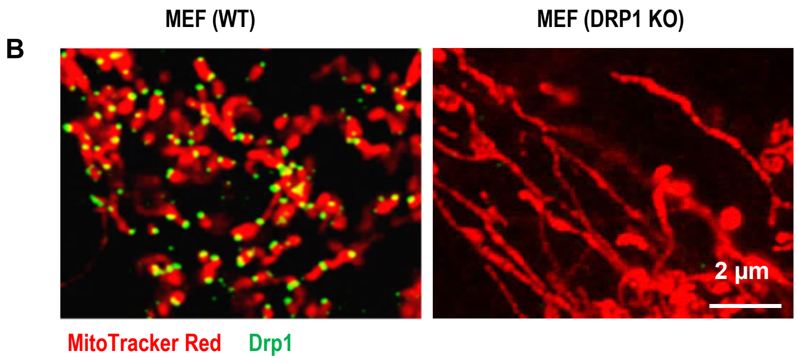

**Fig. S1B. Verification of DRP1 antibody in MEF cells.** Anti-DRP1 antibody (BD Biosciences, 611112) detects mitochondrial associated punctate DRP1 “hot spots” in wild type but not DRP1 KO MEFs. Note that in DRP1 KO cells, mitochondria are dramatically elongated.

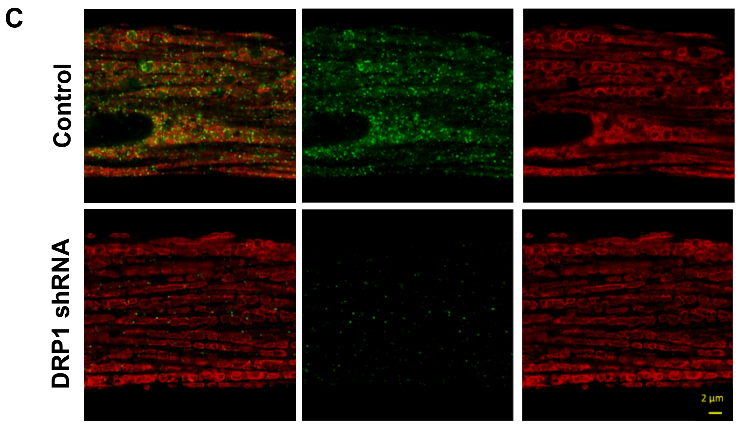

**Fig. S1C. Verification of DRP1 antibody (BD Biosciences, 611112) in freshly isolated cardiomyocytes from rat.**

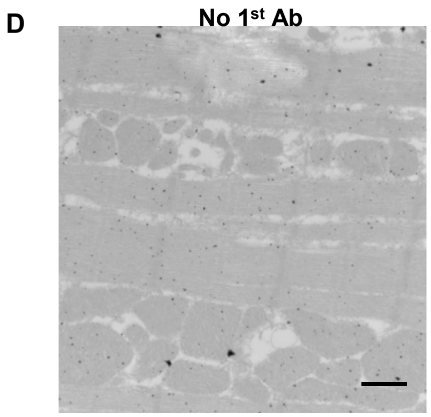

**Fig. S1D. Verification of DRP1 IG secondary antibody signal.**

**Supplementary figure 2**

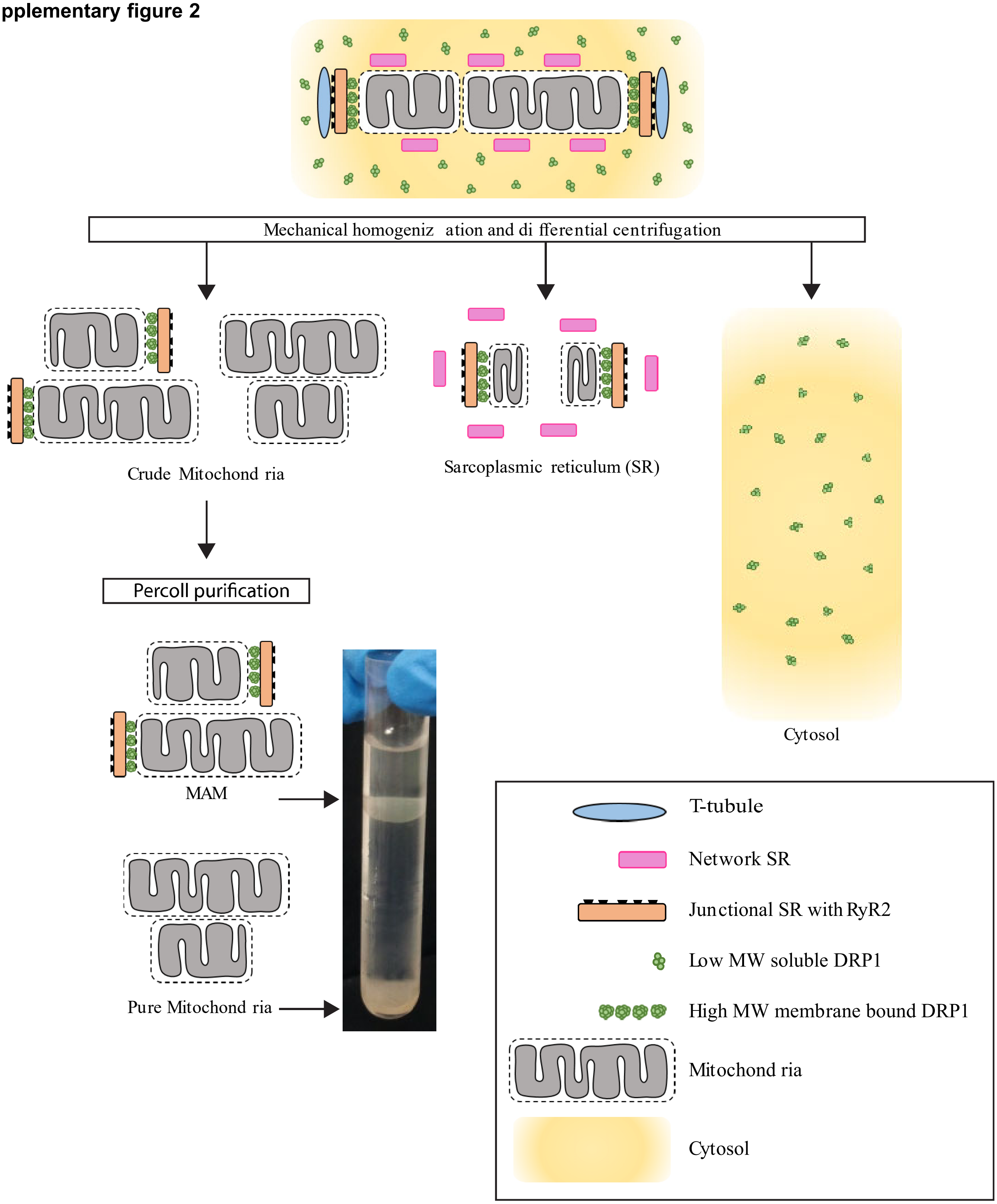

Fig S3. Scheme showing the different components of the cardiac subcellular fractions obtained by differential centrifugation and Percoll gradient purification.

Supplementary figure 3

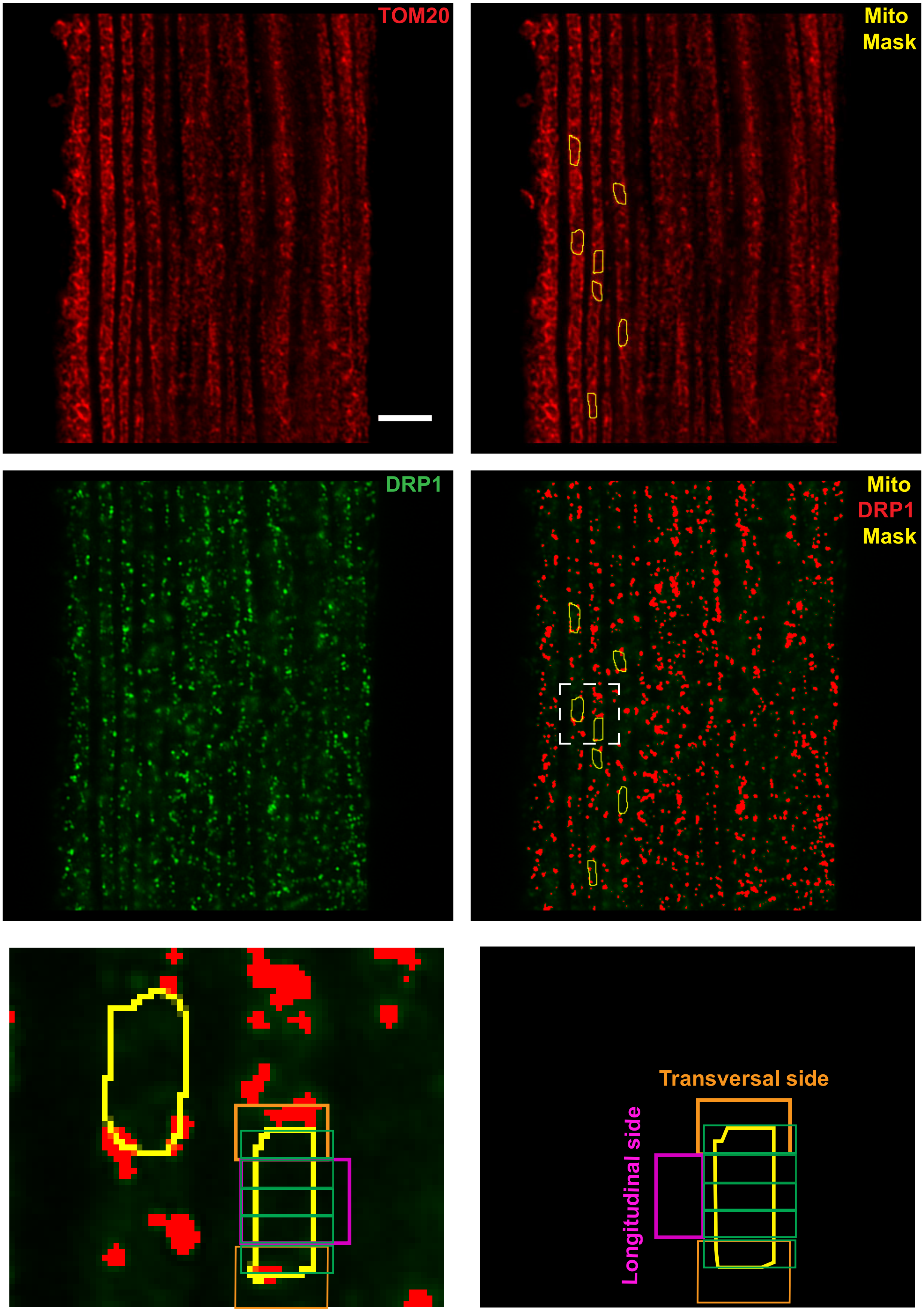

**Fig. S3.** Confocal images of 4x zoom cardiomyocytes regions (scale bar 0.5mm) showing the mitochondrial pattern (TOM20, red, upper left panel) and DRP1 pattern (green, middle left panel). Selected mitochondria for study are labeled in yellow (upper panel, right) and overlapped with the DRP1 mask (5% max intensity threshold). In the lower panels, in higher magnification, the spatial resolution and how each mitochondrion was divided into 5 different regions. The DRP1 thresholded particles touching the mitochondria OMM at the three areas in the middle were considered at the longitudinal side, while the ones touching the mitochondria OMM at the 2 areas in the mitochondrial poles were considered at the transversal side.

---

**Supplementary figure 4**

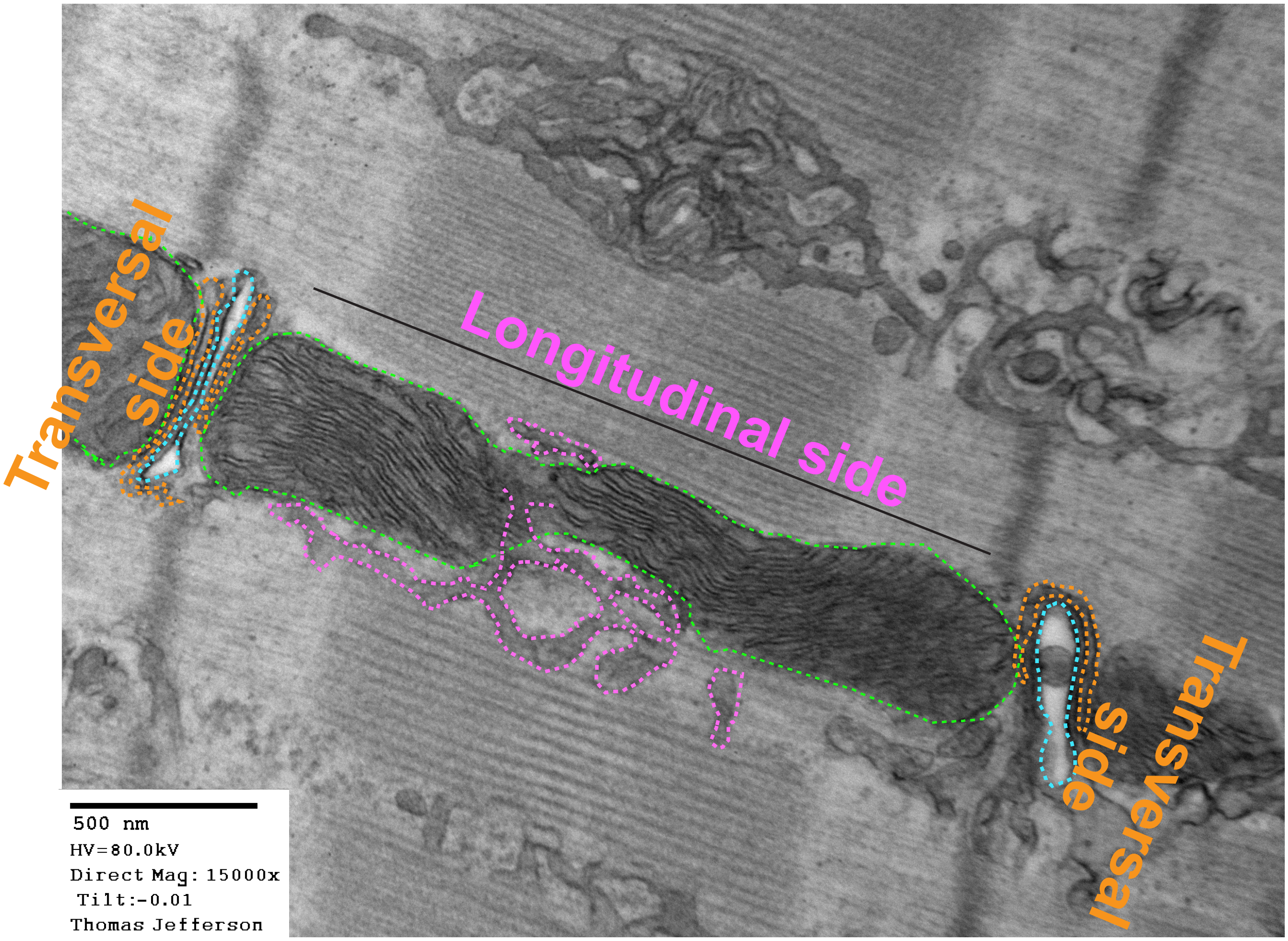

**Fig. S4.** Electron microscopy micrograph of left ventricular papilary muscle showing cardiac interfibrillar mitochondria (green), T-tubules (blue) junctional sarcoplasmic reticulum at the transversal side (orange) of the mitochondria ,and network sarcoplasmic reticulum marking the longitudinal side (pink) of the mitochondria.

**Supplementary figure 5**

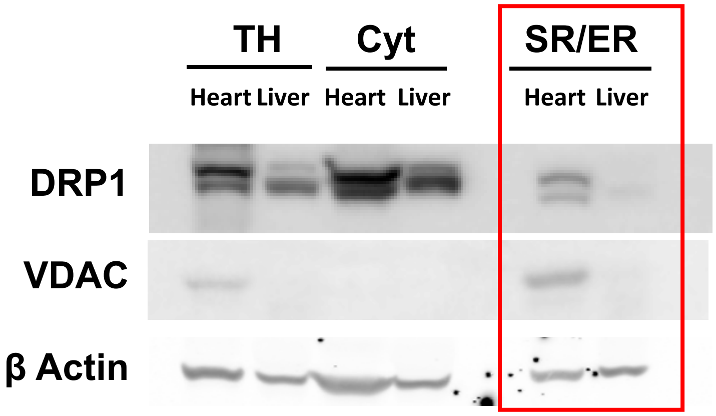

**Fig. S5.** Comparison of DRP1 distribution among SR fractions from hear and Liver. Note that the mitochondria (VDAC) are not detected in the Liver ER fraction in comparison with the high abundance of mitochondria in this fraction in the heart.

**Supplementary figure 6**

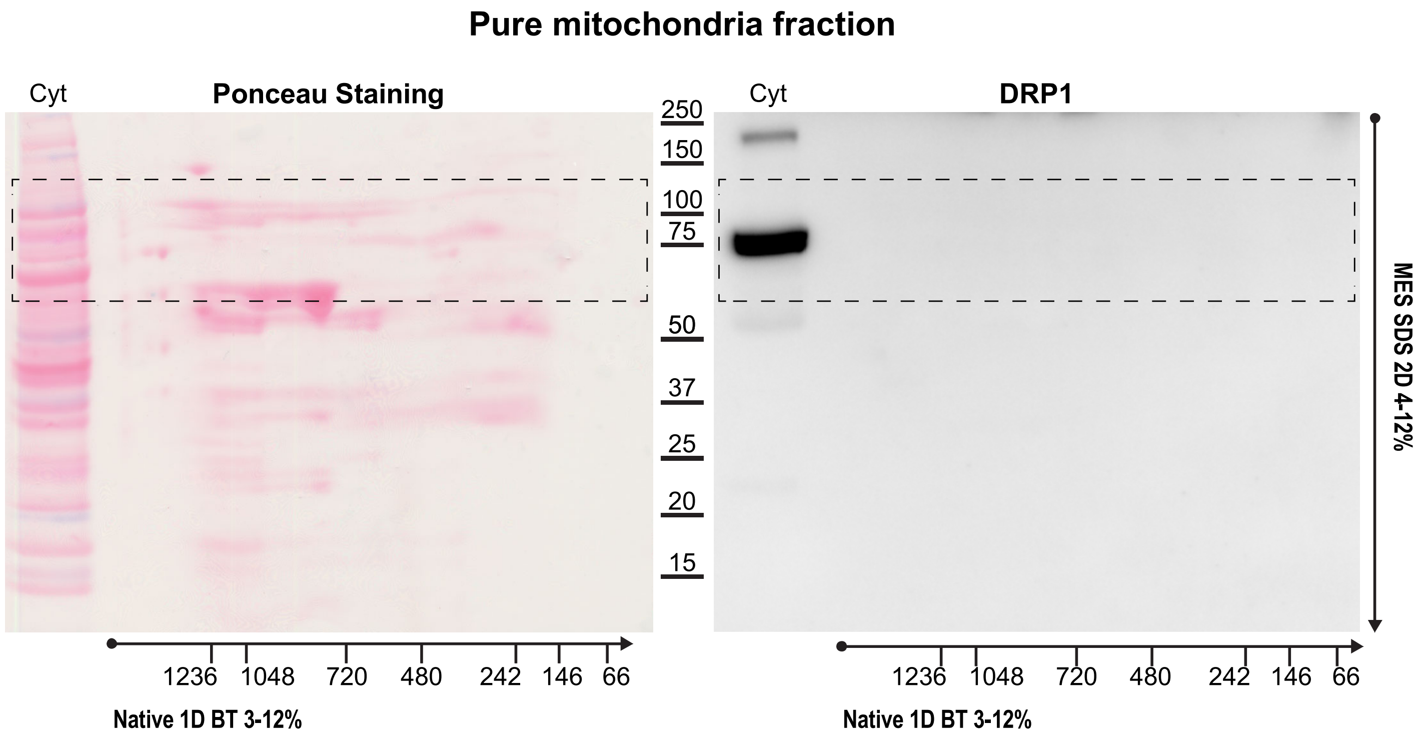

**Fig. S6.** Solubilized proteins from the cytosol, MAM and SR subcellular fractions were separated according to their masses on a linear 3%-12% acrylamide gradient gel for BN-PAGE (first dimension, 1D). Native protein complexes were separated by 4%-12% Bis-Tris SDS-PAGE gradient gel (second dimension, 2D).

**Supplementary figure 7**

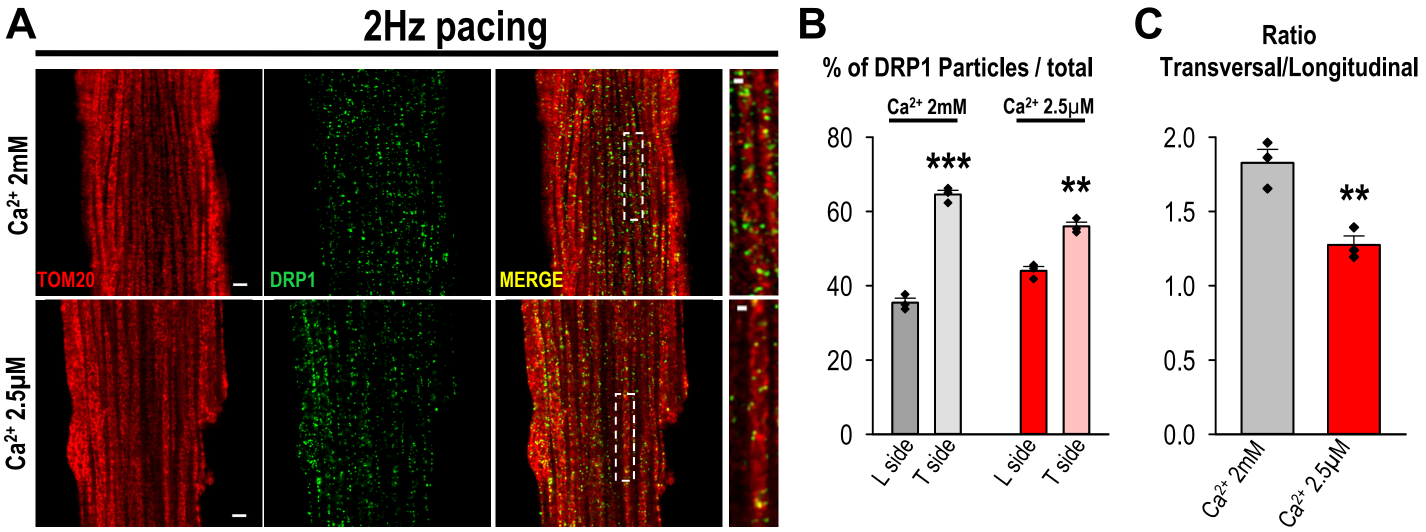

**Fig. S7.** Cardiomyocytes were freshly isolated and submitted to electrical field stimulation (15 min-2 Hz) while being superfused Ca2+ (2 mM) or quasi-Ca2+-free (2.5 μM) buffer. A) Zoomed images (scale bar = 2μ) and detail (scale bar = 0.5μm) are after AiryScan processing DRP1 (green) and TOM20 (red). B-C) Analysis of DRP1 punctate distribution along the mitochondria transversal (T) and longitudinal (L) sides. N = (3 animals; 10 cells per animal) **: 0.01 ≥ p > 0.001, ***:p ≤ 0.001.

**Supplementary figure 8**

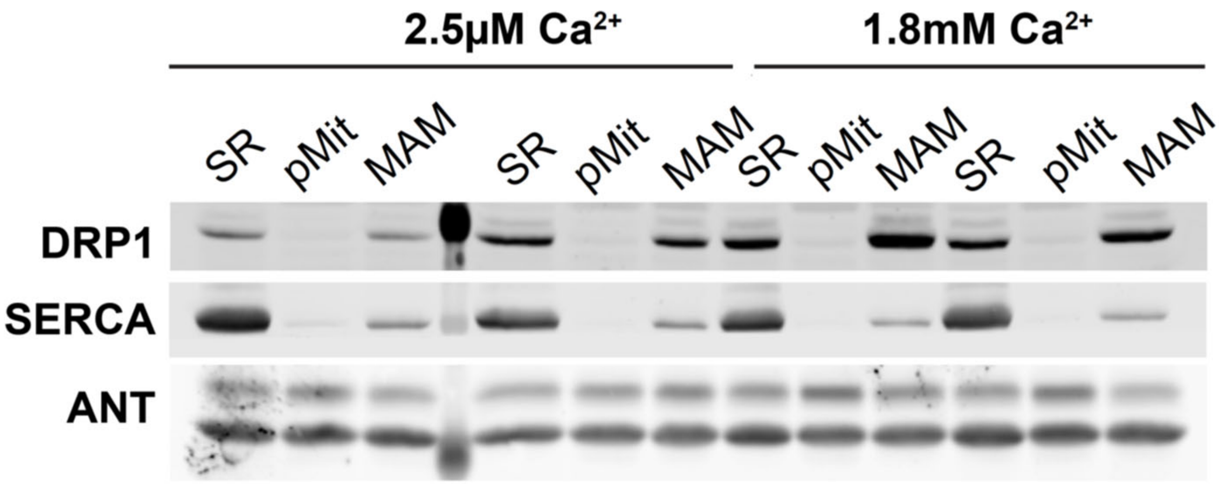

**Fig. S8.** In rat hearts perfused with 2.5μM $Ca^{2+}$ or 1.8mM $Ca^{2+}$ the DRP1 decrease in SR and MAM fractions was not related with DRP1 presence in the pMito fraction.

### Supplementary figure 9

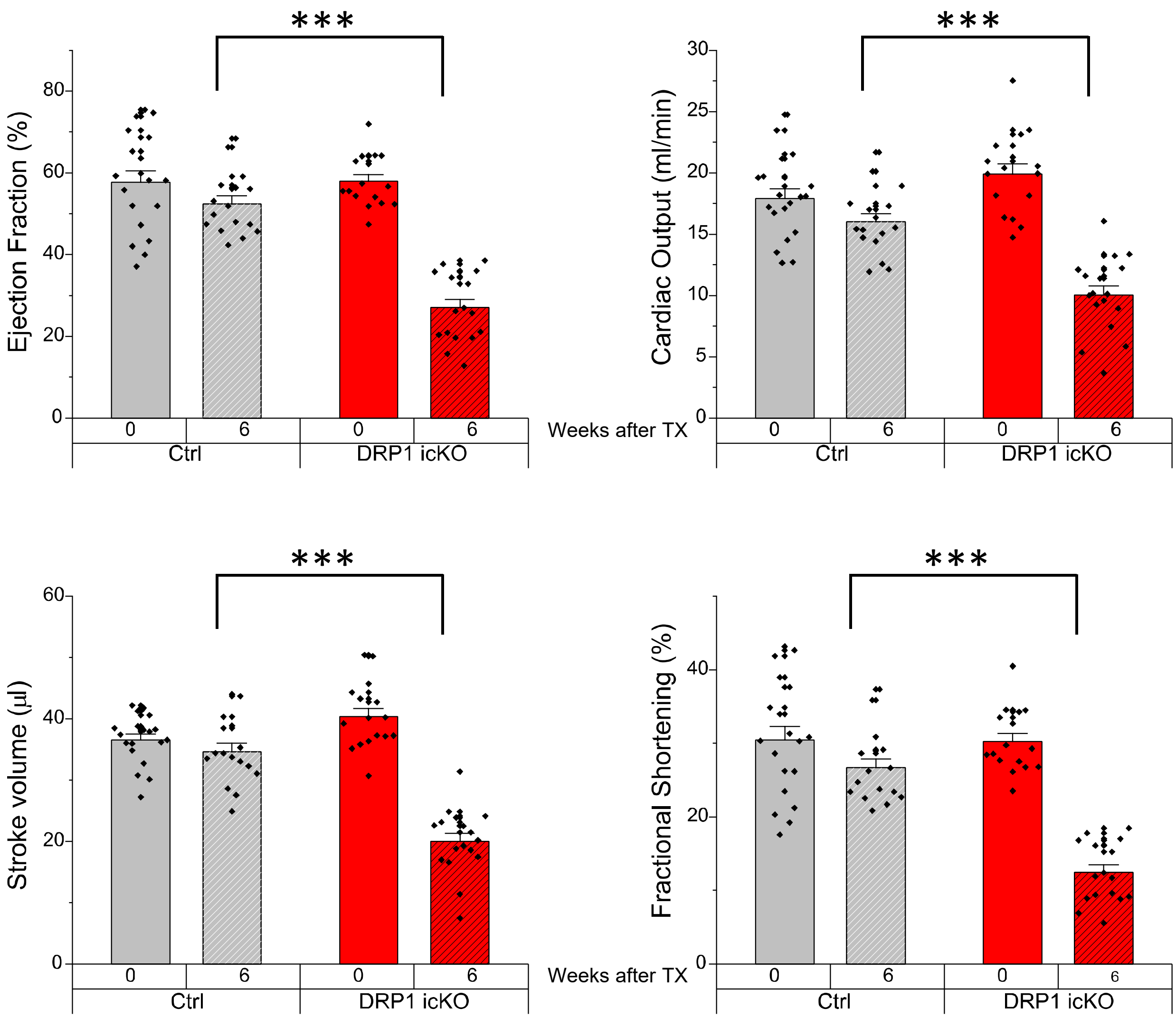

**Fig. S9.** Echocardiographic study of control. animals and DRP1icKO animals before and after 6weeks tamoxifen injection (ctrl N=19, DRP1iCKO N=16).

**Supplementary figure 10**

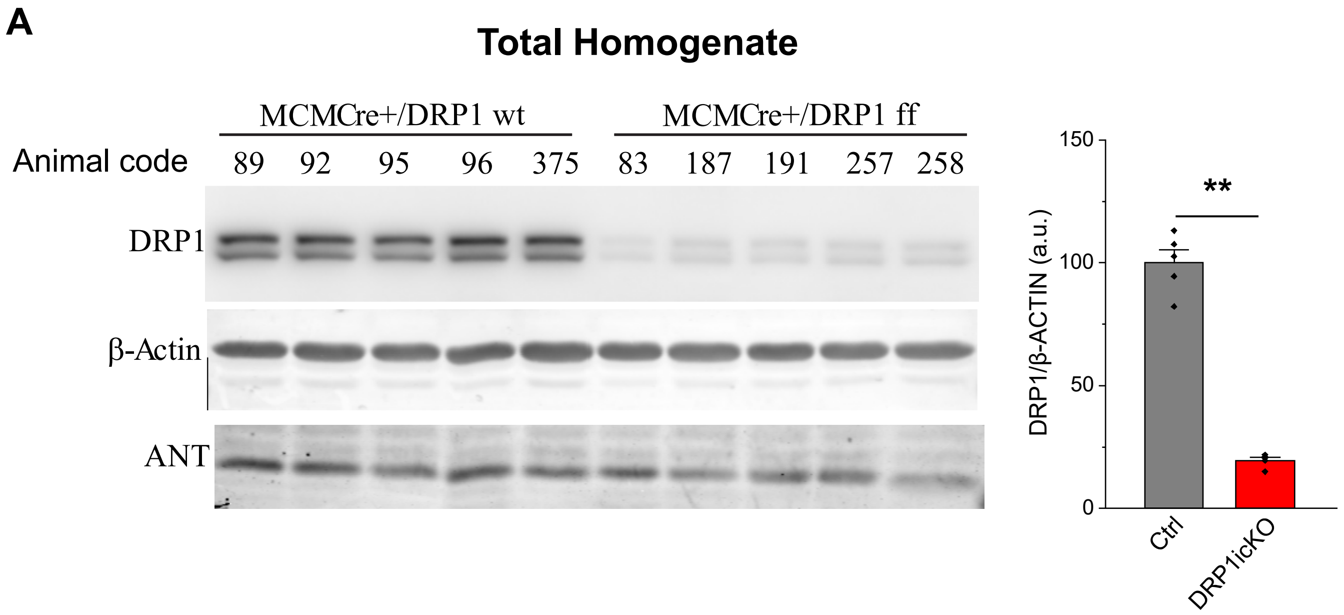

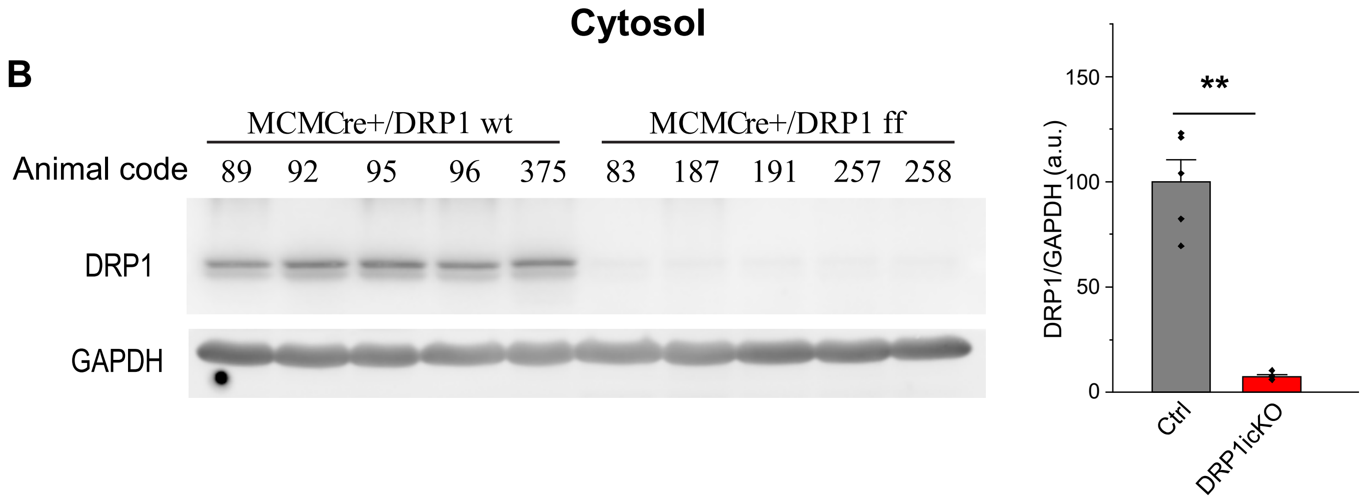

**Fig. S5**. WB analysis and quantification of DRP1 levels of the control and Drp1icko animals in **A)** Total homogenate and **B)** Cytosolic fraction (each number corresponds to a single mice, N=5 per group).

**Supplementary figure 11**

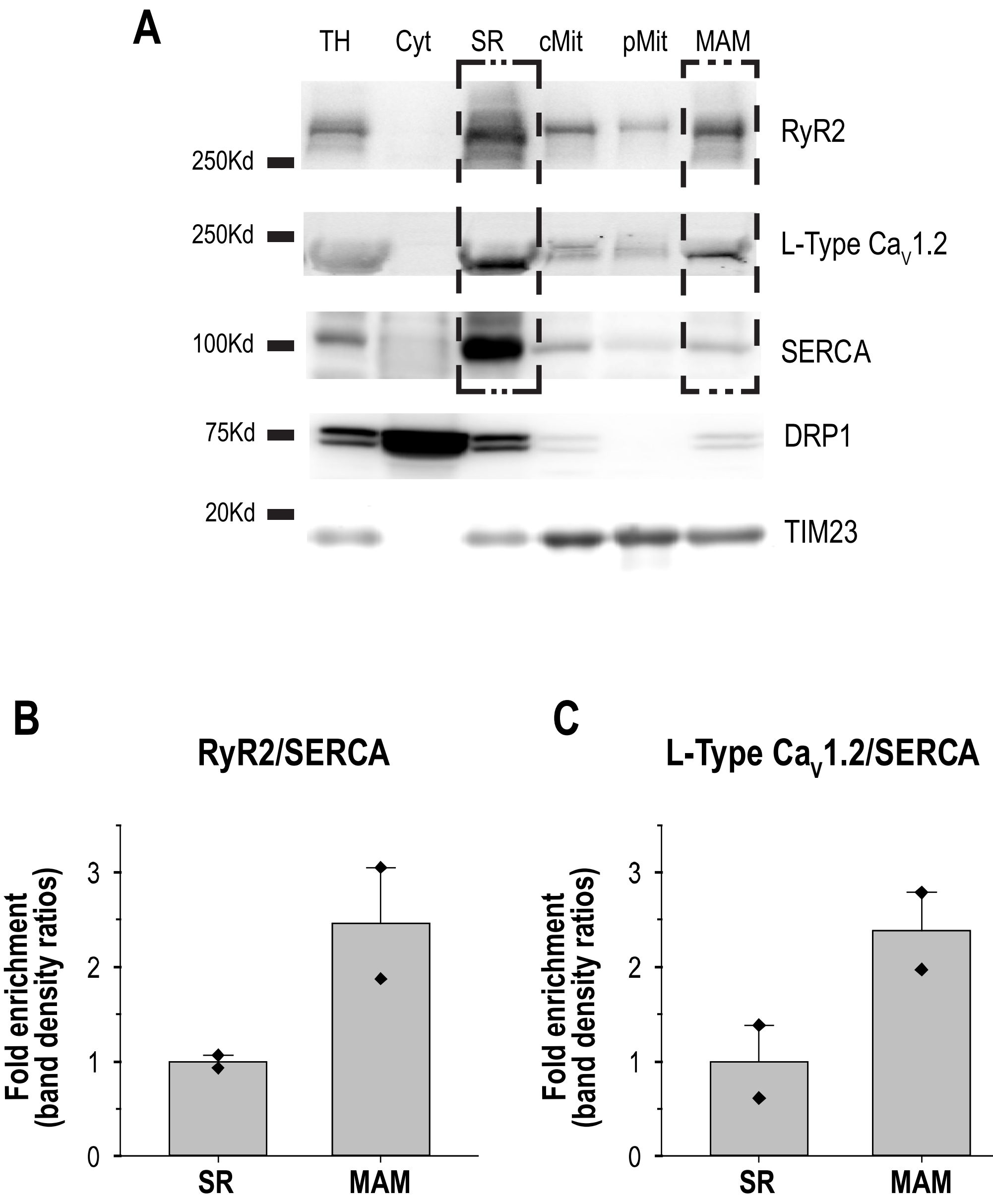

**Fig. S11. A)** Representative Western blot of total cardiac homogenate (TH) and cardiac cellular fractions obtained by differential centrifugation, cytosol (Cyt), sarcoplasmic reticulum (SR), crude mitochondria (cMit) pure mitochondria (pMit) and mitochondria-associated membranes (MAM). RyR2 and L-type Ca2+ channels were used as junctional SR markers while SERCA was used as network SR marker. **B-C)** Comparison of relative abundance of junctional SR and network SR proteins between SR and MAM fractions. N = 3 fractionations from 5 mice hearts each.

### Supplementary figure 12

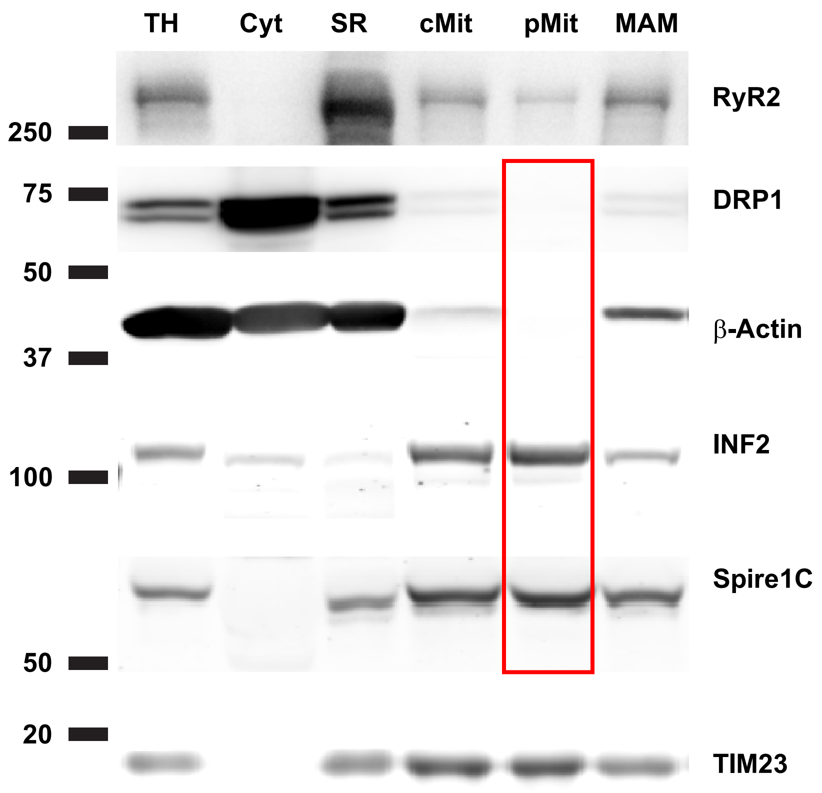

**Fig. S12.** None of the newly described DRP1 related proteins INF2 and Spire1c follow DRP1 distribution among the cardiac cellular fractions. As described before, β-Acting shares same pattern as DRP1 in the different fractions.

**Supplementary table 1. Cytosolic transient parameters at 1Hz**

| | | Ctr | | | DRP1icKO | | | |
|---|---|---|---|---|---|---|---|---|
| | | **Avg** | | **SE** | **Avg** | | **SE** | |
| Ca levels (µM) | Diastolic $Ca^{2+}$ | **0.12** | ± | 0.01 | **0.16** | ± | 0.01 | *p<0.05 |
| | Systolic $Ca^{2+}$ | **0.37** | ± | 0.07 | **0.54** | ± | 0.06 | n.s. |
| Response | Peak time (ms) | **32.65** | ± | 0.57 | **48.55** | ± | 4.26 | **p< 0.01 |
| | tt bl 10% (ms) | **8.53** | ± | 0.24 | **11.49** | ± | 1.46 | n.s. |
| | tt peak 50% (ms) | **16.12** | ± | 0.35 | **20.01** | ± | 1.06 | **p< 0.01 |
| | tt peak 90% (ms) | **24.29** | ± | 0.53 | **32.77** | ± | 1.97 | **p< 0.01 |
| | dep v (µM/s) | **24.09** | ± | 4.62 | **29.43** | ± | 4.20 | n.s. |
| | dep v t (ms) | **16.06** | ± | 0.39 | **19.25** | ± | 1.15 | *p<0.05 |
| Recovery | tt bl 10% (ms) | **51.56** | ± | 2.83 | **81.68** | ± | 6.24 | ***p< 0.001 |
| | tt bl 50% (ms) | **122.18** | ± | 4.35 | **197.78** | ± | 21.25 | **p< 0.01 |
| | tt bl 90% (ms) | **450.50** | ± | 4.10 | **510.10** | ± | 44.00 | n.s. |
| | ret v (µM/s) | **-3.32** | ± | 0.87 | **-3.55** | ± | 0.47 | n.s. |
| | ret v t (ms) | **73.06** | ± | 7.59 | **106.51** | ± | 8.53 | **p< 0.01 |

**Supplementary table 2. Cytosolic transient parameters 5 Hz**

| | | Ctr | | | DRP1icKO | | | |
|---|---|---|---|---|---|---|---|---|
| | | **Avg** | | **SE** | **Avg** | | **SE** | |
| Ca levels (µM) | Diastolic $Ca^{2+}$ | **0.28** | ± | 0.01 | **0.28** | ± | 0.03 | n.s. |
| | Systolic $Ca^{2+}$ | **1.05** | ± | 0.17 | **0.59** | ± | 0.09 | *<0.05 |
| Response | Peak time (ms) | **36.33** | ± | 1.02 | **51.41** | ± | 2.87 | ***p< 0.001 |
| | tt peak 10% (ms) | **11.00** | | 0.62 | **12.88** | | 0.74 | *<0.05 |
| | tt peak 50% (ms) | **19.64** | ± | 0.94 | **24.53** | ± | 1.19 | **p≤ 0.01 |
| | tt peak 90% (ms) | **27.71** | ± | 1.05 | **37.76** | ± | 2.13 | ***p< 0.001 |
| | dep v (µm/s) | **48.85** | ± | 9.70 | **14.02** | ± | 4.06 | **p≤ 0.01 |
| | dep v t (ms) | **19.40** | ± | 1.02 | **23.06** | ± | 1.09 | *<0.05 |
| Recovery | tt bl 10% (ms) | **49.27** | ± | 1.54 | **69.94** | ± | 3.81 | ***p< 0.001 |
| | tt bl 50% (ms) | **85.33** | ± | 2.56 | **113.18** | ± | 4.36 | ***p< 0.001 |
| | tt bl 90% (ms) | **148.07** | ± | 2.56 | **174.06** | ± | 5.41 | **p≤ 0.01 |
| | ret v (µm/s) | **-10.93** | ± | 2.97 | **-3.31** | ± | 0.91 | *<0.05 |
| | ret v t (ms) | **64.27** | ± | 5.67 | **93.35** | ± | 5.73 | ***p< 0.001 |

**Supplementary table 3. Sarcomere length and contractility parameters 1HZ**

| | | Ctr | | | DRP1icKO | | | |
|---|---|---|---|---|---|---|---|---|
| | | Avg | | SE | Avg | | SE | |
| Sarc. Length (µm) | Diastole | **1.772** | ± | 0.015 | **1.698** | ± | 0.016 | **$p < 0.01$ |
| | Amplitude contrac. | **0.16** | ± | 0.02 | **0.09** | ± | 0.01 | **$p < 0.05$ |
| Response | Peak time (ms) | **93.06** | ± | 2.78 | **100.93** | ± | 5.71 | n.s. |
| | tt peak 10% (ms) | **21.69** | ± | 6.71 | **25.98** | ± | 6.29 | n.s. |
| | tt peak 50% (ms) | **28.81** | ± | 0.90 | **42.33** | ± | 2.35 | ***$p < 0.001$ |
| | tt peak 90% (ms) | **59.50** | ± | 1.53 | **73.83** | ± | 3.99 | **$p < 0.01$ |
| | dep v (µm/s) | **-3.88** | ± | 0.62 | **-1.91** | ± | 0.22 | **$p < 0.01$ |
| | dep v t (ms) | **20.75** | ± | 0.84 | **35.18** | ± | 2.57 | ***$p < 0.001$ |
| Recovery | tt bl 10% (ms) | **124.81** | ± | 4.31 | **130.80** | ± | 8.02 | n.s. |
| | tt bl 50% (ms) | **166.13** | ± | 6.29 | **203.75** | ± | 22.61 | n.s. |
| | tt bl 90% (ms) | **307.00** | ± | 27.10 | **494.94** | ± | 58.86 | **$p < 0.05$ |
| | ret v (µm/s) | **2.05** | ± | 0.42 | **1.13** | ± | 0.18 | **$p < 0.05$ |
| | ret v t (ms) | **150.88** | ± | 5.67 | **0.15** | ± | 0.01 | n.s. |

**Supplementary table 4. Sarcomere length and contractility parameters 5Hz**

| | | Ctr | | | DRP1icKO | | | |
|---|---|---|---|---|---|---|---|---|
| | | Avg | | SE | Avg | | SE | |
| Sarc. Length (µm) | Diastole | **1.705** | ± | 0.02 | **1.670** | ± | 0.02 | n.s. |
| | Amplitude contrac. | **0.154** | ± | 0.02 | **0.034** | ± | 0.01 | ***$p < 0.001$ |
| Response | Peak time (ms) | **74.60** | ± | 2.37 | **88.417** | ± | 4.67 | **$p \leq 0.01$ |
| | tt peak 10% (ms) | **19.933** | ± | 1.35 | **34.917** | ± | 3.14 | ***$p < 0.000$ |
| | tt peak 50% (ms) | **36.200** | ± | 2.03 | **52.500** | ± | 3.43 | ***$p < 0.001$ |
| | tt peak 90% (ms) | **56.133** | ± | 2.25 | **73.083** | ± | 3.97 | ***$p < 0.001$ |
| | dep v (µm/s) | **-4.287** | ± | 0.47 | **-0.837** | ± | 0.22 | ***$p < 0.001$ |
| | dep v t (ms) | **33.267** | ± | 2.63 | **49.917** | ± | 3.59 | ***$p < 0.001$ |
| Recovery | tt bl 10% (ms) | **90.857** | ± | 3.20 | **103.000** | ± | 5.36 | n.s. |
| | tt bl 50% (ms) | **111.067** | ± | 3.75 | **127.417** | ± | 5.43 | *$<0.05$ |
| | tt bl 90% (ms) | **142.067** | ± | 4.66 | **171.909** | ± | 6.22 | **$p \leq 0.01$ |
| | ret v (µm/s) | **3.418** | ± | 0.41 | **0.698** | ± | 0.20 | ***$p < 0.001$ |
| | ret v t (ms) | **106.000** | ± | 3.94 | **118.667** | ± | 6.14 | n.s. |
